# Population coding of predator imminence in the hypothalamus

**DOI:** 10.1101/2024.08.12.607651

**Authors:** Kathy Y.M. Cheung, Aditya Nair, Ling-yun Li, Mikhail G. Shapiro, David J. Anderson

**Affiliations:** Division of Biology and Biological Engineering, California Institute of Technology; Pasadena, CA, USA; Tianqiao and Chrissy Chen Institute for Neuroscience Caltech; Pasadena, CA USA; Department of Neurobiology, School of Basic Medical Sciences, Capital Medical University, Beijing, China; Howard Hughes Medical Institute; Chevy Chase, MD, USA; Andrew and Peggy Cherng Department of Medical Engineering, California Institute of Technology; Pasadena, CA USA; Division of Chemistry and Chemical Engineering, California Institute of Technology; Pasadena, CA, USA

## Abstract

Hypothalamic VMHdm^SF1^ neurons are activated by predator cues and are necessary and sufficient for instinctive defensive responses. However, such data do not distinguish which features of a predator encounter are encoded by VMHdm^SF1^ neural activity. To address this issue, we imaged VMHdm^SF1^ neurons at single-cell resolution in freely behaving mice exposed to a natural predator in varying contexts. Our results reveal that VMHdm^SF1^ neurons do not represent different defensive behaviors, but rather encode predator identity and multiple predator-evoked internal states, including threat-evoked fear/anxiety; neophobia or arousal; predator imminence; and safety. Notably, threat and safety are encoded bi-directionally by anti-correlated subpopulations. Finally, individual differences in predator defensiveness are correlated with differences in VMHdm^SF1^ response dynamics. Thus, different threat-related internal state variables are encoded by distinct neuronal subpopulations within a genetically defined, anatomically restricted hypothalamic cell class.

**Highlights:**

1. Distinct subsets of VMHdm^SF1^ neurons encode multiple predator-evoked internal states.
2. Anti-correlated subsets encode safety vs. threat in a bi-directional manner
3. A population code for predator imminence is identified using a novel assay
4. VMHdm^SF1^ dynamics correlate with individual variation in predator defensiveness.

## Introduction

Across the animal kingdom, rapid and effective defensive behaviors are critical to protect an animal from being harmed or killed by predators or conspecifics. Predator threats induce specific innate defensive responses that do not require observational or reinforcement learning. Studies in rodents have provided evidence that innate and learned (conditional) defensive responses are mediated by anatomically distinct neural pathways (Figure 1A)^1–4^.

**Figure 1.**
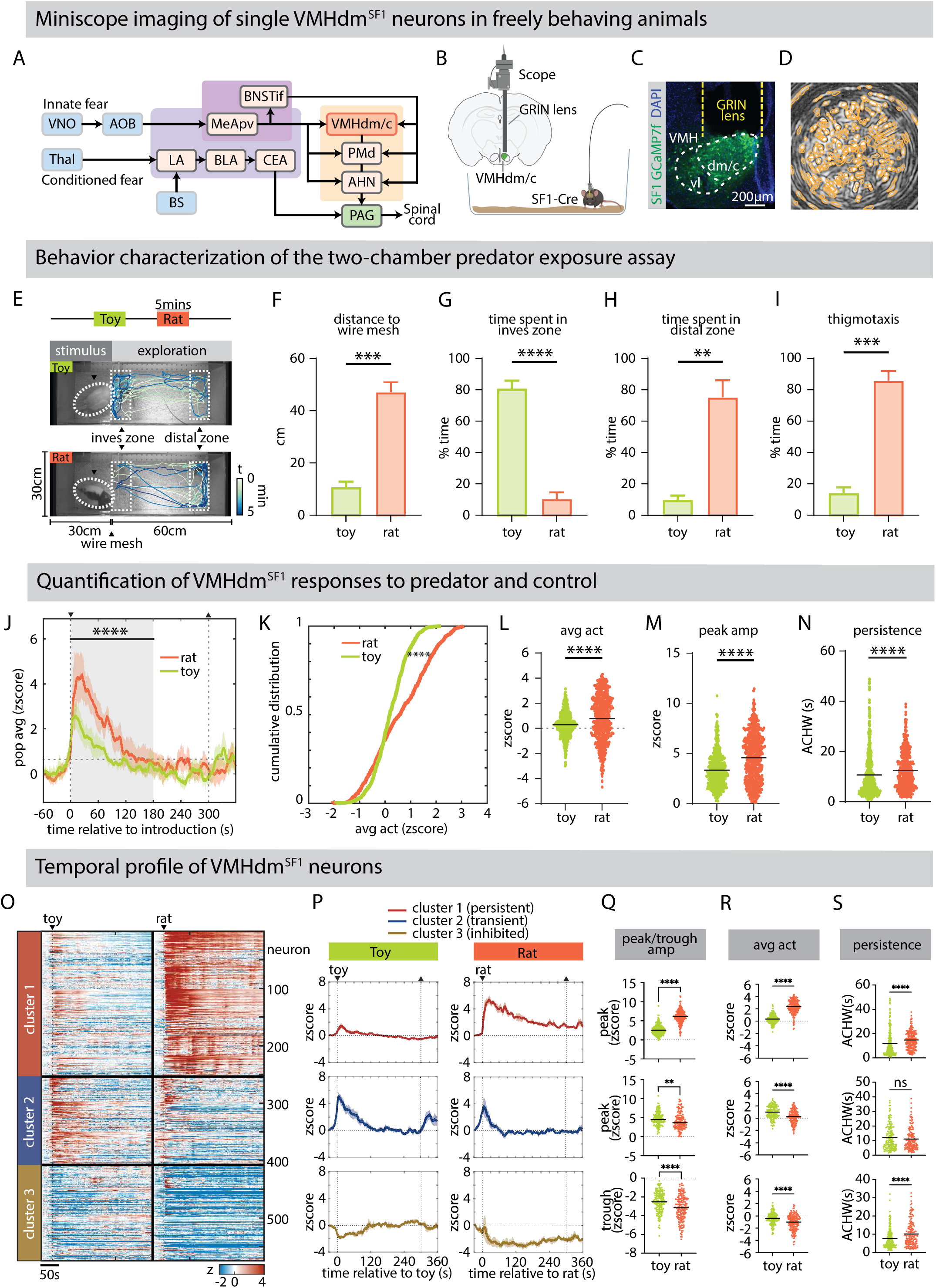
Diverse temporal patterns of single VMHdm^SF1^ neurons in freely behaving animals. (A) Schematic summarizing circuits for innate and conditioned fear. (B) Microendoscopic imaging of VMHdm in a freely behaving SF1-Cre mouse. (C) GCaMP7f expression in VMHdm^SF1^ neurons. Green: SF1-positive neurons with GCaMP7f expression; Blue: DAPI staining. (D) Field of view of an imaged mouse with GCaMP7f expression in VMHdm^SF1^ neurons. (E) Design of the two-chamber predator exposure arena and schematics of the assay overlaid with the trajectory of a representative animal (blue traces). The rat/toy is behind a wire mesh barrier. (F-I) Graphs showing (F) average distance to wire mesh, (G) % time spent in the inves zone, (H) % time spent in distal zone, and (I) thigmotaxis during toy and rat presentations. \*\**P*<0.01, \*\*\**P*<0.001,\*\*\*\**P*<0.0001. Inves: investigation. (n = 5 mice) (J) Mean ± s.e.m. population calcium responses of VMHdm^SF1^ neurons during rat (red) and toy (green) presentations, aligned to stimulus introduction. Shaded area indicates the window used for quantification in K-N. \*\*\*\**P*<0.0001. (K) Cumulative distribution of VMHdm^SF1^ single cell average activity during rat and toy. \*\*\*\**P*<0.0001. (n = 574 cells, 5 mice) (L-N) (L) Average activity, (M) peak amplitude, and (N) ACHW of VMHdm^SF1^ single cell activity during rat and toy. \*\*\*\**P*<0.0001. ACHW: autocorrelation halfwidth; peak amp: peak amplitude; avg act: population average activity. (O) Cell raster and k-means clustering of neural dynamics during rat and toy presentation. Red: cluster 1; blue: cluster 2; brown: cluster 3. (P) Mean ± s.e.m. population responses of VMHdm^SF1^ single cells in each cluster during rat and toy presentations. (Q-S) (Q) Peak/trough amplitude, (R) average activity, and (S) ACHW of VMHdm^SF1^ single cell activity in each cluster during rat and toy presentations. ns: not significant, \*\**P*<0.01, \*\*\*\**P*<0.0001. See also Figures S1, S7.

The dorsomedial and central subdivisions of the ventromedial hypothalamus (VMHdm/c) play a pivotal role in the mammalian innate defensive network^3,5–13^. This region receives information from olfactory/pheromonal auditory and other sensory processing areas that are activated by threat-related stimuli^3^. It then integrates and relays this information to downstream structures to drive defensive behaviors^5,6,11,14–19^.

Glutamatergic neurons in VMHdm/c that express the nuclear receptor protein steroidogenic factor 1 (SF1; also known as NR5a1), hereafter referred to as VMHdm^SF^^1^ neurons, are necessary for predator-evoked defensive behaviors in mice^6–8,20^. Moreover, optogenetic stimulation of these neurons has a negative valence, and can elicit different defensive behaviors in a threshold-dependent manner^6^. Different defensive behaviors can also be triggered by activating different projections from VMHdm, such as those targeting the anterior hypothalamic nucleus (AHN) or periaqueductal gray (PAG)^13^. In addition, previous neural recordings performed in VMHdm have confirmed that these neurons are strongly and persistently activated during a predator encounter^10–12^.

Although these studies have demonstrated VMHdm’s necessity and sufficiency for mediating defensive behaviors^6,13^, as well as their activation by predator-derived cues^7,9–12^, it is still unclear what variables are being represented by the level of activity or the temporal dynamics of VMHdm^SF1^ neurons during a naturalistic predator encounter. Previous studies have left open the question of whether VMHdm^SF1^ neurons mainly encode the identity of a threatening stimulus^9,10^, or other features of the stimulus such as its intensity, modality and/or duration of exposure^10^; pre-motor programs for defensive behaviors; or an internal affective state of defensiveness or fear.

Earlier work has shown that the VMHdm^SF1^ population is activated during naturalistic threat exposure and that different subsets of VMHdm^SF1^ neurons represent different predator odors or auditory cues^9–12^. Electrophysiological recordings of anonymous units in VMHdm have revealed distinct subpopulations whose activity is correlated with either threat investigation or subsequent flight, respectively^12^. Calcium imaging of single VMHdm^SF1^ neurons in head-fixed animals has revealed that predator-evoked responses decay slowly over tens of seconds following stimulus removal^10^. This persistent activity has been shown to be required for persistent post-threat defensive behavior, such as thigmotaxis, suggesting that VMHdm^SF1^ neurons encode an internal defensive state that endures beyond its inciting stimulus^10,21^. However, it remains unclear whether the duration of persistent activity encodes the duration of the defensive response, or whether the level of activity encodes the intensity or type of response.

Understanding what features of a predator encounter are encoded or represented in the pattern of VMHdm^SF1^ activity is challenging. Because the stimulus is innately threatening, it is difficult to distinguish whether the activated neurons represent threat object identity versus a defensive internal state evoked by the threat (or both). In other brain regions, threat (or negative valence) is represented independently of stimulus identity^22,23^. In studies of conditional fear in the amygdala, the distinction between these two stimulus-linked features is more easily dissociated^24^. Prior to conditioning, the stimulus is neutral; hence its meaning to the animal as a threat or threat-predicting signal is only acquired after conditioning. Therefore, any changes in the neural response pattern evoked by the stimulus after conditioning likely reflect its acquired ability to induce fear and promote defensive behavior, rather than its identity *per se*. In contrast, innately threatening stimuli evoke internal defensive states and motor responses upon the animal’s first encounter.

In this study, we have investigated neural coding by individual VMHdm^SF1^ cells in freely behaving mice during naturalistic predator encounters. Using computational methods, we have attempted to identify the variables represented by this activity, at both the single-cell and population levels. While the stimulus itself accounts for a large fraction of variance in VMHdm^SF1^ neural activity, little variance was explained by motor behavior, suggesting that the residual variance might encode an internal state variable^25–27^. Consistent with this interpretation, we identified discrete subpopulations of VMHdm^SF1^ neurons that encode threat vs. safety bi-directionally, as well as a population-level representation of “predator imminence”^28–34^. We also found a strong positive correlation between individual differences in predator defensiveness and the duration of persistence of predator-evoked activity. These data, taken together with previous studies^6,10,11,13^, provide evidence that VMHdm^SF1^ neurons encode and control the strength and length of a predator-evoked internal affective state, which may contribute to the subjective experience of fear in humans^4,21,33,34^.

## Results

### Heterogeneous temporal profiles of VMHdm^SF1^ neurons

We performed microendoscopic imaging in VMHdm^SF1^ neurons expressing GCaMP7f in freely behaving mice from a stereotaxically injected Cre-dependent AAV vector (Figures 1B-D)^10,35–37^. Initial behavioral experiments were performed in a custom-designed two-chamber arena, which consisted of a stimulus compartment and an exploration chamber that were separated by a wire mesh (Figure 1E). For measurement purposes, we defined two zones: an investigation (“inves”) zone near the rat; and a “distal” zone at the opposite end of the cage. Male mice were first acclimated to the exploration chamber for 10 minutes followed by 5 minutes of baseline recording, and then exposed sequentially to two different stimuli: a control (toy rat) followed by a live rat, each presented for 5 minutes separated by a 5-minute interval. The order of stimulus presentation was fixed to avoid habituation to the rat and consequent suppression of responses to the control. The introduction of the rat caused a distinct alteration in the animals’ spatial navigation patterns in comparison to the control. The animals stayed further away from the stimulus chamber, spent more time in the distal zone, avoided the inves zone and exhibited an increase in thigmotaxis (Figures 1F-I).

We collected imaging data from a total of 600 neurons in 5 mice. Live rat presentation evoked a stronger and more persistent population average response compared to the toy rat control (Figures 1J, S1A-C). Across individual cells, the rat also evoked significantly higher average activity (p<0.0001) and peak amplitude (p<0.0001) than the control (Figures 1K-M). Moreover, individual cells also exhibited longer persistence (p<0.0001) in response to the rat than to the toy, measured using autocorrelation half-width^36,38,39^ (Figures 1N, S1D). Most toy-activated cells overlapped with those activated by the rat, while approximately half of the imaged cells were activated by both rat and the toy (Figure S1E). However, we identified subpopulations that were selectively tuned to either rat or toy (Figure S1F-H), consistent with previous studies that showed VMHdm^SF1^ encodes stimuli identities^9,10^.

To reduce the dimensionality of the dataset, we next used unsupervised k-means clustering to classify neurons by their correlated temporal profiles during rat presentations, with the cluster number set at 3 (Figure 1O). This resulted in three clusters that exhibited distinct average responses to toy vs. rat presentation (Figures 1P-S). Rat presentation evoked strong and persistent activation in cluster 1 but relatively little activity during toy presentation; we refer to this for convenience as the “persistent cluster”. In contrast, cluster 2 exhibited strong but relatively transient activation to both rat and toy; we refer to this as the “transient cluster”. In contrast to the previous two clusters, cluster 3 was strongly and persistently inhibited by the rat as well as by the toy, but to a much lesser extent; we refer to this as the “inhibited cluster”. Although it has been reported previously that a subset of VMHdm^SF1^ neurons is inhibited by rat, their function is uncharacterized^10,12^. Thus our unsupervised k-means clustering revealed distinct subpopulations of VMHdm^SF1^ neurons, each exhibiting diverse and distinct responses to the rat and toy.

### Relationship between VMHdm^SF1^ neural activity and predator proximity or motor activity

We next examined the dynamics of VMHdm^SF1^ activity during live rat presentation. The mice frequently approached and inspected the rat in the investigation zone and subsequently fled from the threat to the distal zone (Figure 2A). This behavioral sequence, referred to as “risk assessment”, is commonly performed to evaluate and respond to a threatening situation^30^. We separated the risk assessment sequence into approach and escape phases and examined the associated neural dynamics. Close investigation of the rat evoked a rise to a peak in VMHdm^SF1^ population average activity, followed by a decline as the animal fled (Figure 2B, black line). This observation prompted us to investigate whether this rise and fall in VMHdm^SF1^ activity simply represents changes in predator proximity, reflecting the distance-dependent intensity of predator odors^25^, or rather in the level of perceived threat evoked by the predator.

**Figure 2.**
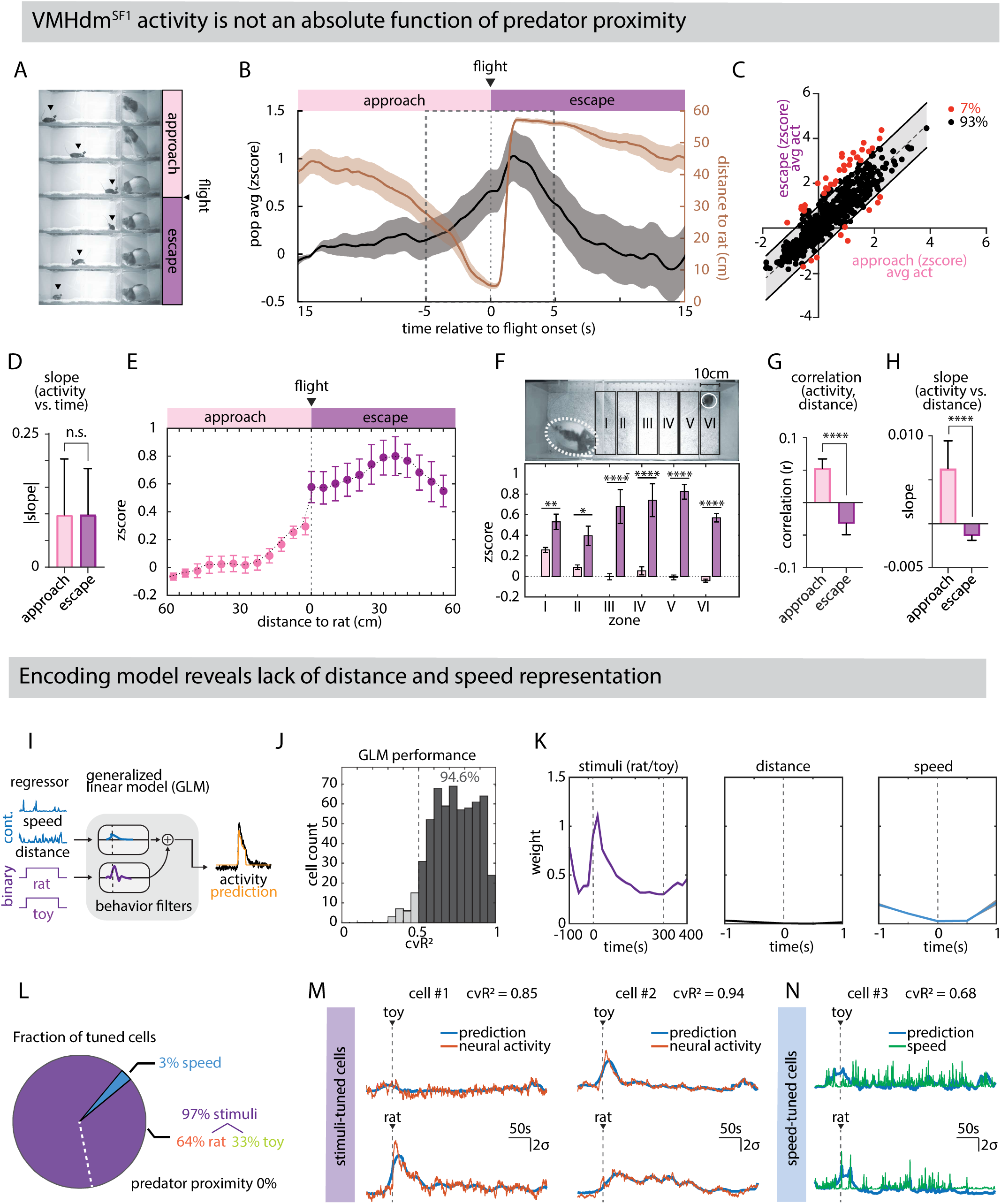
VMHdm activity is not a direct measure of predator proximity or motor intensity. (A) Time series of video stills illustrating typical sequence of mouse behaviors following rat stimulus introduction, separated into approach (pink) and escape (purple) phases by initiation of flight. Black arrows indicate the position of the mouse. (B) Mean ± s.e.m. population responses of VMHdm^SF1^ cells during rat investigation (black), overlaid with distance to rat (brown), aligned to the onset of flight. Dashed line indicates the window used for quantification in C-H. (n = 5 mice) (C) Relationship between the average activity of VMHdm^SF1^ single cells during approach and escape. Black and red points, inside and outside 95% confidence interval, respectively. (n = 574 cells, 5 mice) (D) Mean slope of VMHdm^SF1^ single cell activity vs. time during approach and escape phases. ns: not significant. (E) Mean ± s.e.m. response of VMHdm^SF1^ neurons as a function of distance to the rat during approach and escape phases. (F) Activity is not a function of absolute distance to rat. Upper: Activity was binned into 6 virtual zones. The width of each zone is 10 cm. Lower: Neural activity of VMHdm^SF1^ neurons during approach (pink bars) and escape (purple bars) phases in each zone. *P<0.05, **P<0.01,****P<0.0001. (G) Mean Pearson’s correlation coefficients between the VMHdm^SF1^ single cell activity and distance to rat during approach and escape phases. ****P<0.0001. (H) Mean slope of VMHdm^SF1^ single cell activity vs. distance during approach and escape phases. ****P<0.0001. (I) GLM architecture for detecting cells that were selectively tuned to speed, distance to rat, the presence of rat and the presence of toy, along with behavioral filters. A non-negativity constraint was applied aid the interpretation of GLM weights. (J) GLM performance across all cells analyzed (n = 574 cells, 5 mice). (K) Kernel weights for each regressor. (L) Fractions of cells tuned to each regressor. (M-N) Example cells tuned to (M) stimuli and (N) speed. See also Figure S1.

To resolve this ambiguity, we examined changes in VMHdm^SF1^ neuronal activity within a time window spanning 5 sec before and 5 sec after flight onset (Figure 2B, dashed vertical lines). We then plotted the average activity of each neuron during the former (approach) interval against its average activity during the latter (escape) interval (Figure 2C). Most of the neurons (93%) were distributed along the diagonal, indicating that each cell exhibits a similar average activity level during approach vs escape. The absolute rate of change in average neural activity over time within this window was also similar during approach and escape (albeit opposite in sign), suggesting that it represents a common process or variable (Figure 2D).

We observed that during approach vs. escape, the population average activity differed over the same distance range from the rat (0-60 cm; Figure 2E). To quantify this difference, we divided the arena into 10cm bins, namely zone I (closest to rat) to zone VI (furthest from rat; Figure 2F, upper). In all zones, neural activity was significantly higher during escape than during approach (Figure 2F, lower). This and other parametric differences (Figures 2G-H) suggest that VMHdm^SF1^ activity level is not simply a function of absolute predator proximity. During approach, VMHdm^SF1^ activity may in part reflect rising sensory cue intensity (Figure S1I). However, given its continuously high activity level during the escape phase, even at the maximum distance between the predator and the animal (Figures 2E-F, S1J), a more parsimonious explanation is that the signal encodes a variable that is integrated during approach and which decays during escape with a similar time constant, perhaps encoding an internal defensive state evoked by the rat. This would account for the similar (but opposite-direction) rates of change in activity as a function of time during both phases (Figures 2B, D).

This interpretation, however, did not exclude that high activity during the escape phase reflects the vigorous motor activity associated with flight, rather than a defensive internal state (Figure S1K). To rigorously disambiguate these alternatives, we employed a generalized linear model (GLM) to identify neurons that are selectively tuned to the stimuli (rat or toy), predator proximity, and the speed of the animal (Figure 2I)^40^. The GLM was able to fit 94.6% of the cells with good performance (cvR^2^>0.5; Figure 2J). Among all the regressors, the stimuli (rat/toy) contributed most of the kernel weight to the model fit, while velocity and distance accounted for little to no weights, respectively (Figure 2K). This principled approach confirmed that the majority (97%) of the well-fit VMHdm^SF1^ neurons are primarily modulated by the stimuli (rat/toy) (Figure 2L-M). Among the 97% stimuli-tuned cells, 64% of the neurons’ GLM fit had a higher weight contributed by the rat than by the toy and the remaining 33% had the opposite. None of the cells was found to be predator proximity-tuned (Figure 2L). 3% of the well-fit neurons were found to be tuned to the speed of the animal (Figures 2L, N). Since the number of these speed-tuned neurons is low, and can potentially reflect motion artifacts, we decided not to investigate further this subpopulation.

Thus, our GLM analysis revealed that stimulus identity (rat or toy) was the primary source of variance in VMHdm^SF1^ neural activity (Figure 2L). However, this observation does not distinguish whether VMHdm^SF1^ neurons simply encode the identity of the stimulus object, or rather encode an internal state(s) evoked by the different stimuli. Furthermore, the total variance in VMHdm neural activity was not fully explained by absolute distance to the predator or motor patterns, suggesting the residual variance might encode an internal state variable as well (Figure S1L)^25–27^.

### Distinct VMHdm^SF1^ subpopulations exhibit opposite responses to threat

To investigate further the relationship between stimulus identity and internal state coding, we designed a behavioral assay in which the animal’s internal state can be manipulated independently of the stimulus object’s identity^41^. We modified the two-chamber arena by introducing a shelter (refugium) in the middle of the exploration chamber (Figure 3A, dashed black outline). Under these conditions, the animals robustly altered their spatial navigation pattern after encountering the rat, spending more time in the shelter and distal zone than during toy presentation (Figures 3B-D). Unlike in the two-chamber assay without a refugium (Figures 1E-I), the animals did not exhibit a statistically significant increase in thigmotaxis during rat vs toy exposure, although there was a trend (Figure 3E). This could in part reflect a switch in defensive strategy in response to the change in context – i.e., the availability of a refugium.

**Figure 3.**
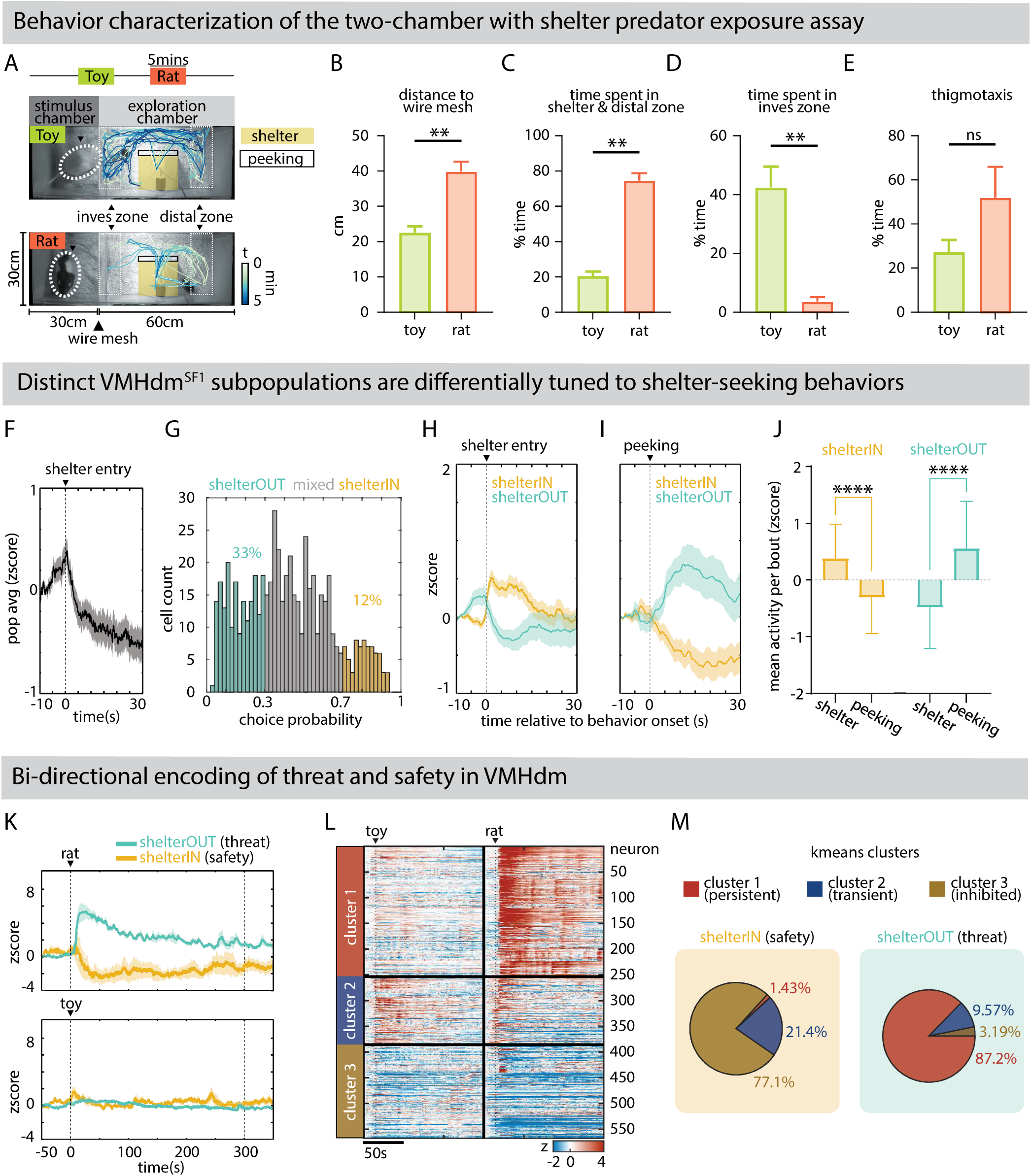
VMHdm has distinct subpopulations that represent safety and threat signals. (A) Illustration of the two-chamber predator exposure arena with shelter overlaid with the trajectory of a representative animal (blue traces). Yellow shaded area: shelter; black outline: peeking zone. (B-E) Graphs showing (B) average distance to wire mesh, (C) % time spent in the shelter and distal zone, (D) % time spent in inves zone, and (E) thigmotaxis during toy and rat presentations. Inves: Investigation. (n = 5 mice) (F) Mean ± s.e.m. population responses of VMHdm^SF1^ neurons during shelter-seeking behavior, aligned to shelter entry. (n = 5 mice) (G) Choice probability histogram and percentages of shelter-tuned units. ShelterOUT (teal): 33%, ShelterIN (amber): 12%. (n = 567 cells, 5 mice) (H-I) Mean ± s.e.m. population responses of shelterIN and shelterOUT cells, aligned to (H) shelter entry and (I) peeking. (J) Responses of shelterIN (left) and shelterOUT (right) cells during shelter and peeking. \*\*\*\**P*<0.0001. (K) Mean ± s.e.m. population responses of shelterIN and shelterOUT cells across entire interaction time following presentations of the rat (upper) and toy (lower). (L) Cell raster and unsupervised k-means clustering of neural dynamics during rat and toy presentation. (M) Pie charts showing the composition of ShelterIN (left) and ShelterOUT (right) cell populations by k-means clusters. See also Figures S2, S7.

Under these conditions, we observed an overall reduction in population average activity that was time-locked to shelter entry (Figure 3F). However, this observation did not distinguish whether this reduction reflected a change in the overall activity of a homogeneous cell population, or rather changes in the relative activity of different subpopulations of VMHdm^SF1^ neurons. To dissect the neural representation of the shelter entry response with single-cell resolution, we applied choice probability analysis to VMHdm^SF1^ single-unit activity to identify cells that are selectively modulated by shelter-seeking behaviors (Figure 3G)^42–44^. Unexpectedly, the results revealed two distinct subpopulations with opposite responses, one whose activity was higher inside (“ShelterIN” cells), and the other outside, the shelter (“ShelterOUT” cells; Figures 3H, S2A-B). These two subpopulations exhibited time-locked and opposite responses upon shelter entry. Prior to exiting the shelter, mice often cautiously extended their heads out of the shelter, presumably to assess their surroundings. We refer to this behavior as “peeking”. During peeking behavior, we observed a rapid switch in neural responses: ShelterIN neurons became inhibited, and ShelterOUT neurons were activated, which was opposite to that observed upon shelter entry (Figures 3I-J).

To our surprise, these two subpopulations also exhibited opposite responses at the time of introduction of the live rat stimulus. ShelterOUT neurons showed strong activation upon the introduction of the live rat, while ShelterIN neurons were strongly inhibited (Figure 3K, upper). Furthermore, these responses were long-lasting: the two subpopulations remained well-separated throughout the entire rat presentation. These differential dynamics were not observed during toy presentation (Figure 3K, lower). Taken together, these results indicate that the response of VMHdm^SF1^ neurons to a threatening object can be modified by context (specifically the presence of a shelter). This in turn suggests that VMHdm^SF1^ may encode an internal state of perceived threat or “fear” that is flexibly modulated, in addition to (or instead of) the identity or proximity of a threatening object.

We attempted to investigate the relationship between the ShelterIN and ShelterOUT subpopulations and the k-means clusters identified previously in the two-chamber assay without the shelter (Figures 1O-S). Because it was not possible to track individual cells across these two assays, we performed an independent unsupervised k-means clustering on the dataset from the assay with the shelter, with the number of clusters set to 3 as well. This resulted in clusters that exhibited qualitative similarities to the 3 clusters obtained from the assay without the shelter (Figures 1O-S, 3L, S2C-G). The majority of the ShelterIN neurons (77.7%) were assigned to cluster 3 (inhibited) while 87.2% of the ShelterOUT subpopulation were in cluster 1 (persistent; Figure 3M).

### Sparse motor representation of defensive behaviors in VMHdm^SF1^

Previous research has demonstrated that optogenetic stimulation of VMHdm^SF1^ neurons induces various types of defensive behaviors in an intensity or projection-dependent manner^6,13^. According to the Predator Imminence theory, animals adjust their defensive strategies — such as nesting, avoidance, freezing, flight, and panic-like behaviors (jumping, undirected escape and defensive attack)—depending on whether and when they expect to encounter a predator (Figure 4A)^28,29,31,32^. ‘Imminence’ reflects the prey’s perceived likelihood of being attacked, or predicted time to contact with the predator^28,29^. In humans, verbal reports of predator-evoked fear are inversely related to predator distance^33,34^. However, the perceived likelihood of attack is also influenced not only by predator proximity but also by other factors, for instance, the physical context of the surrounding environment (such as the availability of an escape route or escapability i.e. the amount of space available)^28,29,31,32,45–47^. To explore the encoding of defensive behaviors, predator imminence and context by VMHdm^SF1^ neural activity, we developed the Predator Imminence and Context Assay (PICA), a novel behavioral paradigm inspired by this theoretical framework (Figures 4A-B)^28,29,48^.

**Figure 4.**
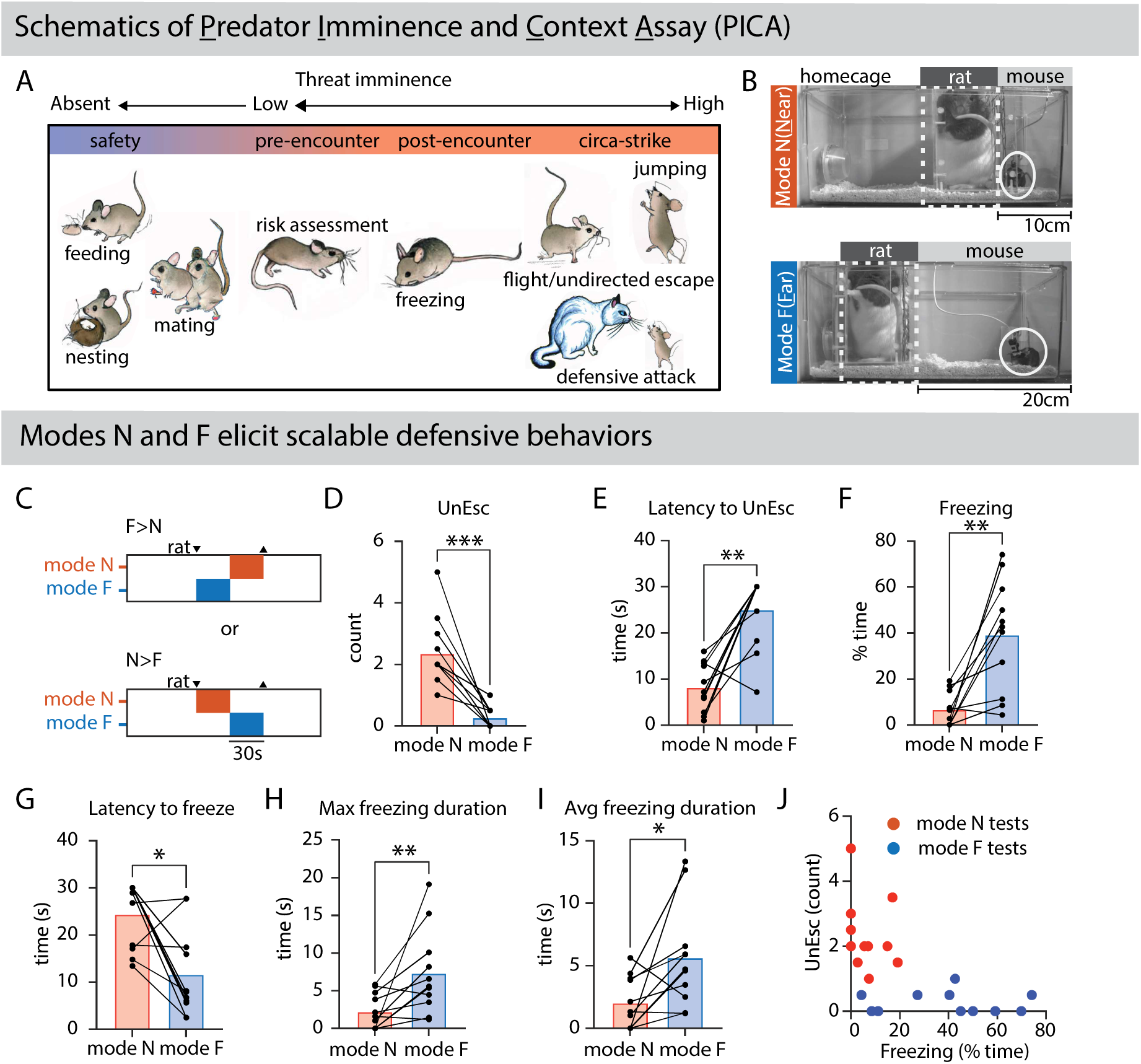
A novel behavioral paradigm to examine the encoding of defensive behaviors, predator imminence, and contexts. (A) Illustration of different defensive modes across the predator imminence continuum (based on a model developed by Fanselow & Lester, 1988^28^). Rodent-specific defensive behaviors are described for each mode. The theory describes that an escalating fear level is associated with increasing intensity of defensive behaviors. (B) Illustration of the predator imminence and context assay (PICA). Dashed white line: rat enclosure; solid white circle: mouse; relative size of spaces occupied by rat and mouse are indicated above images. (C) Schematics and presentation sequence in the PICA assay. Red: mode N; Blue: mode F. (D-I) Comparison of the (D) number of undirected escapes (UnEsc), (E) latency to undirected escape, (F) % time spent freezing, (G) latency to freeze, (H) maximum freezing duration and (I) average freezing duration in modes N and F. Connected data points represent individual animals. \**P* >0.05, \*\**P*<0.01, \*\*\**P*<0.001. (n = 11 mice) (J) Summary of relative frequencies of freezing vs. undirected escape behavior displayed during modes N vs. F animals. All animals were implanted with miniscopes. For data on wild-type non-surgical animals see Figure S3A-E.

We used PICA to simulate a naturalistic predator encounter and evoke a wider spectrum of innate defensive behaviors, for instance, jumping, freezing and undirected escape, which were not or rarely observed in the two-chamber assay. We also wished to achieve better temporal and spatial control over the distance between the mouse and the predator, in comparison to the two-chamber setup. For technical reasons, we were not able to continuously vary the distance between the mouse and a live predator in a controlled manner^49^. Instead, PICA has two distinct conditions that differ in predator distance and available escape space, each designed to provoke specific defensive behaviors from the mouse^28,29,33,34,47,48^. In this protocol, the rat was presented in a transparent plastic enclosure (Figure 4B, white dashed outline) within the mouse’s home cage. The rat’s container, matching the width of the mouse’s home cage, was sealed on all sides to prevent leakage of rat urine into the mouse’s home cage except it was perforated on the side facing the mouse to facilitate odor transfer while preventing direct contact. The PICA featured two modes, mode N (<u>n</u>ear) and mode F (<u>f</u>ar), distinguished by the position of the rat enclosure which was under the experimenter’s control. All animals were subjected to a standardized 60s exposure to the rat that comprised 30s in mode N and 30s in mode F, with the sequence of presentation systematically counterbalanced (Figure 4C).

In mode N, the rat enclosure was positioned ∼10cm away from the cage wall with the mouse trapped in between, reducing the mouse’s available space to approximately one-third of the original home cage’s maximum area (Figure 4B, upper). High-intensity defensive behaviors, such as undirected escape and jumping, were frequently observed in this mode, while freezing was minimal (Figures 4D-J, S3A-C). Conversely, in mode F, the rat enclosure was positioned further back, against the opposite cage wall (∼20cm away), expanding the mouse’s available movement space to two-thirds of the total home cage area (Figure 4B, lower). In mode F, animals predominantly engaged in freezing behavior, while undirected escape and jumping were less frequent (Figures 4D-J, S3A-C). Based on prior studies, we conclude from these behavioral results that mode N provokes an active defense strategy, while mode F resembles passive defense^28,29,31,32,45,46,48^. Notably, when the two modes were presented in sequence, the animals rapidly adjusted their qualitatively different defensive strategies to the changing conditions, independent of the order in which the modes were presented (Figures S3F-H).

In mode N, animals with a head-mounted miniscope showed frequent undirected escapes, which were characterized by rapid charging between corners of the home cage, causing bedding materials to pop into midair. However, they showed less or no jumping compared to wild-type non-surgical (WTNS) animals, likely due to the weight (∼2g) and tethering of the miniscope (Figures 4D-E). On the other hand, freezing behavior in both modes was unaffected by the miniscope (Figures S3D-E). Despite the behavioral difference observed in mode N between miniscope-bearing and WTNS animals, the idea that mode N evokes more intense defensive behaviors than mode F still appears valid. Thus, through manipulation of threat context and distance, the PICA assay enabled the reliable elicitation of distinct defensive behaviors under conditions that permitted simultaneous single-cell resolution neural recording in freely behaving animals (Figure 4J).

We next used the PICA assay to search for neural correlates of different types of defensive behaviors within the VMHdm^SF1^ neuron population (Figure S3I). Initially, we hypothesized that different defensive behaviors are controlled by distinct VMHdm^SF1^ subpopulations, similar to the amygdala in mediating conditioned fear^50^. Surprisingly, neither freezing nor undirected escape correlated with discernible, time-locked changes in the overall activity of VMHdm^SF1^ neurons (Figures S3J-L). After accounting for variance contributed by rat presentation, freezing and undirected escape explained less than 10% of the remaining variance in single-unit VMHdm^SF1^ neural activity (Figure S3M). A GLM was applied to rigorously identify single-cell correlates of freezing and undirected escape (Figure S3N)^40^. However, fewer than 1% of cells in the total population analyzed showed GLM weights on these behaviors, while approximately 70% were responsive primarily to rat presentation (Figure S3O).

### A scalable representation of threat context emerges from population coding

Since we did not identify any neural correlate of qualitatively different behaviors in VMHdm^SF1^ neurons, we hypothesized that VMHdm^SF1^ might instead encode quantitative variation in the level of intensity of an internal defensive state that is triggered by the two modes which are associated with distinct defensive strategies^10,28,29,33,34,48^. Naively, one might have expected that as threat imminence increased, overall activity in VMHdm^SF1^ neurons might increase as well (i.e., that mode N has higher activity than mode F). Surprisingly, however, modes N and F evoked similar average activity levels and dynamics in VMHdm^SF1^ neurons when presented independently (Figures 5A-B, S4A-G). (While the majority of VMHdm^SF1^ neurons were activated by the rat presentation, a subset of the neurons was inhibited (Figure S4H-N); the latter could potentially correspond to the safety neurons identified in the two-chamber assay.) When the two modes were presented sequentially— first N then F (N>F) and vice versa (F>N), VMHdm^SF1^ neurons also exhibited similar population average activity level and dynamics in each mode, characterized by a rapid rise to peak during the first mode, followed by slow decay (Figures 5C-D). Because of this slow decay, the second mode presented always exhibited a lower VMHdm^SF1^ response than the first mode, irrespective of the mode phase (Figures S4O-R). This pattern is consistent with our previous observations of VMHdm^SF1^ activity in head-fixed animals, where repeated presentations of the same stimulus yielded rapidly decreasing responses, likely reflecting habituation^10^.

**Figure 5.**
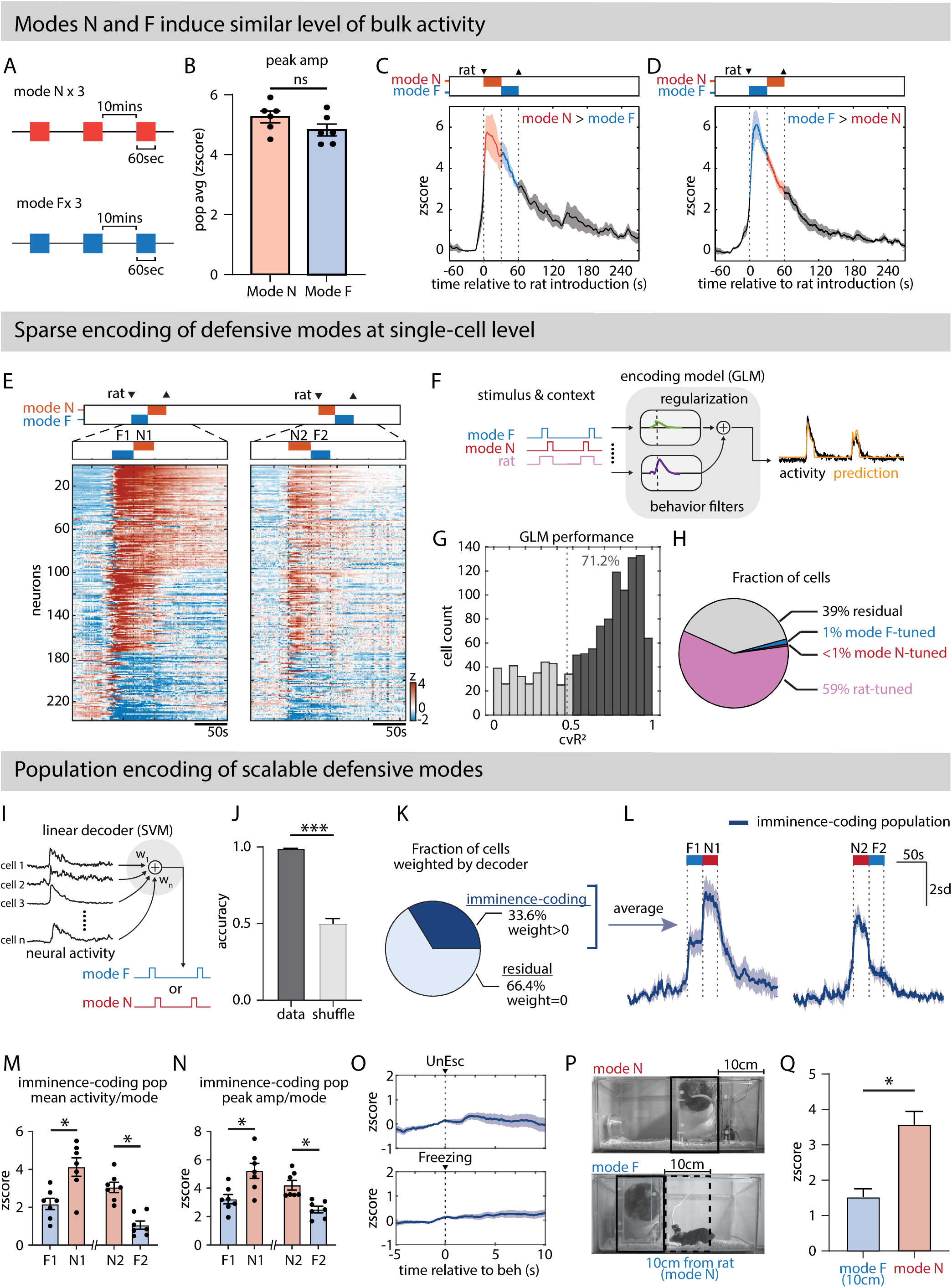
Encoding of defensive states emerges from population mechanisms. (A) Schematics of the behavior paradigm in which each animal was presented with either mode N or F for 60 seconds, repeated three times at 10-minute intervals. The same animal was subjected both mode N and mode F tests, one week apart, with the order systemically balanced. (B) Mean peak amplitude of VMHdm^SF1^ average population activity during modes N and F, averaged across three bouts for each mouse. (n = 6 mice) (C-D) Mean ± s.e.m. population responses of VMHdm^SF1^ neurons during (C) N>F and (D) F>N. (N>F, n = 5 mice, F>N, n = 6 mice). Different groups were used for the two test variants to avoid habituation to the rat stimulus. (E) Schematics of the double-reversed PICA (F1>N1, N2>F2) and representative z-scored neural activity raster plots from an example animal. F1: mode F in the first presentation; N1: mode N in the first presentation; N2: mode N in the second presentation; F2 mode F in the second presentation. (F) GLM architecture for detecting cells that were selectively tuned to each defensive mode. Each neuron was trained using annotations of mode N, mode F, and rat presence (Mode N + Mode F) as regressors, along with behavior filters. A non-negativity constraint was applied to aid the interpretation of GLM weights. (G) GLM performance from 1349 cells across 7 mice. (H) Fraction of cells significantly modulated by each regressor. (I) Non-negative linear decoder (SVM) architecture for predicting mode N vs. mode F, trained on VMHdm^SF1^ single-cell neural activity. (J) Performance of SVM decoder in predicting N vs. F. \*\*\**P*<0.001. (n = 1349 cells, 7 mice) (K) Fraction of cells weighted by the decoder. Cells weighted by the decoder are termed the “imminence-coding population”. (L) Mean ± s.e.m. responses of imminence-coding population over time during PICA tests. (M-N) (M) Average response and (N) peak amplitude of mean activity of imminence-coding population during each mode in PICA. *P<0.05. (O) Mean ± s.e.m. responses of imminence-coding population aligned to undirected escape (upper) and freezing (lower). (P) Dashed outline indicates the region in mode F (10 cm from the rat) in which the predator proximity is the same as in mode N, used to compute metrics in (Q). (Q) Mean activity of the imminence-coding population when the mouse was within 10 cm from the rat during mode N and mode F. \**P*<0.05. See also Figures S3F-O, S4, S5, S6 and S7.

The apparently rapid habituation of VMHdm^SF1^ activity to the predator stimulus within trials consisting of sequential presentation of the two stimuli (N>F or FN) presented a significant challenge in attempting to extract a signal encoding defensive mode per se. To compensate for habituation, we implemented a “double-reversed” PICA, wherein the two defensive modes were sequentially presented, and subsequently, after a 10-minute pause, presented again but in the reverse order (Figure 5E). With this modified PICA, we next addressed the question of whether differences in mode N vs. mode F might be encoded by VMHdm^SF1^ signals that are masked by total population activity and would necessitate the investigation of single-cell activity and dynamics.

We first utilized GLMs in an attempt to identify single-cell correlates of the two defensive modes (Figure 5F). However, GLMs revealed very few cells (∼1%) that were specifically tuned to one or the other defensive modes (Figures 5G-H). The majority of the neurons (59%) were found to be tuned simply to the presence of the rat. However, by training a linear support vector machine (SVM) decoder on single-unit neural activity, it was possible to predict whether the animal was in mode N or mode F with high accuracy (Figures 5I-J). This result suggested that an encoding of defensive modes indeed exists in VMHdm^SF1^ neurons, but at the population rather than the single-cell level.

To gain further insight into how defensive modes are encoded in VMHdm^SF1^, we focused on cells that were weighted on the mode N vs. mode F decoder, hereafter referred to as the “imminence-coding population” (which accounted for 33.6% of the total imaged population), to examine their neural dynamics (Figures 5K, dark blue fraction, S5A). The mean activity of the imminence-coding population scaled with the two modes in a manner that correlated with the level of predator imminence built into each mode^28^: the average activity was higher in mode N than in mode F, regardless of presentation order (Figures 5L-N, S5B-C), in contrast to the monotonic decrease across two sequential modes seen in the residual population (Figures S5D-H). Most strikingly, the observed increase in mode N, in trials where mode N followed mode F, was in the opposite direction to what would be expected for within-trial habituation (Figure 5L, left), as reflected by bulk VMHdm^SF1^ dynamics (Figures 5C-D). Importantly, behavior-triggered average analysis indicated that neither freezing nor undirected escape were associated with any time-locked response in the imminence-coding population (Figure 5O).

To account for the possibility that the higher level of activity in the imminence-coding population in mode N was due simply to a higher level of predator odorant in the more confined space, we examined the period in mode F when the animals voluntarily approached the rat to within the same 10cm distance as allowed in mode N (Figure 5P). Under these conditions, in theory, the animals should have been exposed to the same odorant concentration in both modes during such periods. However, the imminence-coding population still displayed a significantly lower level of activity in mode F at the same distance from the rat as in mode N, even after normalizing for the different times of exposure at this distance in the two modes (p<0.05; Figure 5Q). In addition, the activity of the imminence-coding population did not vary linearly with distance to the rat (Figure S5I). These data suggest that the population activity identified by SVM decoder analysis can act as a neural correlate of predator imminence and/or encounter context.

Subsequently, we attempted to investigate the relationship between the imminence-coding population and the k-means clusters identified in previous experiments (Figures 1O, 3L). We performed an independent k-means clustering on the double-reversed PICA dataset, with the number of clusters set to 3 as well. This again resulted in clusters that exhibited qualitative similarities to the 3 clusters obtained previously (Cluster 1: persistent, Cluster 2: transient, Cluster 3: inhibited; Figure S5J). All three clusters contributed to the imminence-coding population (Figure S5K). However, cluster 1 (persistent) was preferentially weighted by the imminence-coding population (41.5%), cluster 3 (inhibited) ranked second with 36.5% and cluster 2 (transient) contributed the least with only 22%. Since cluster 1 has the highest fraction among the imminence-coding population, we hypothesized that cluster 1 contributed the most to the encoding of predator imminence and/or threat context. We applied the linear SVM decoder on neural activity extracted from cluster 1 neurons only (Figure S5L) and it was still possible to distinguish mode N from mode F with very high accuracy (Figure S5M). Notably, the mean activity (not weighted average) of the cluster 1 neurons weighted by this decoder resembled the dynamics of the imminence-coding population qualitatively, exhibiting a higher level of activity in mode N than mode F regardless of the presentation order (Figure S5N).

### Variation in VMHdm^SF1^ neural dynamics correlates with variation in the level of defensive behavior across individuals

In the PICA assay, we observed a wide range of differences among genetically identical individuals in the amount of defensive behaviors they expressed (Figures 4D-J). Given that overall VMHdm^SF1^ neural activity does not encode defensive behavior *per se* in the PICA assay (Figures S3I-O), we exploited this behavioral variability to investigate whether it was correlated with variability in any features of VMHdm^SF1^ activity, as a way to identify parameters that might encode or represent the intensity of behavioral defensiveness.

To quantify defensiveness, we created a new metric termed the ’defensive index’ (DI), which amalgamates measurements of freezing and undirected escape behaviors, thereby offering a combined assessment of the total defensive behaviors exhibited by each animal in N>F or F>N trials in the PICA assay. The DI comprises two components: 1) the percentage of time spent freezing, and 2) the number of undirected escapes. Since the level of VMHdm^SF1^ neural activity is not correlated with freezing or undirected escape, for the purposes of this comparison we assigned equal weight to both behaviors in calculating the DI for each animal. As the highest number of undirected escapes observed in a single trial was 8, we standardized both factors on a scale from 0 to 10. Undirected escape was quantified as an absolute count. The percentage of time spent freezing, originally ranging from 0 to 100%, was rescaled to a 0-10 range, with 50% freezing equivalently represented as a score of 5. The sum of these two components formed the DI. Importantly, all DI measurements were based on each animal’s first exposure to a live rat in their lifetime, to avoid any potential habituation or learning effects. Naively, one might have expected that animals which displayed more defensive behaviors (higher DI) would have a higher level of VMHdm^SF1^ activity. However, the DI showed no correlation (r^2^ = 0.01) with the peak amplitude of VMHdm^SF1^ total population average activity (Figures 6A, C).

**Figure 6.**
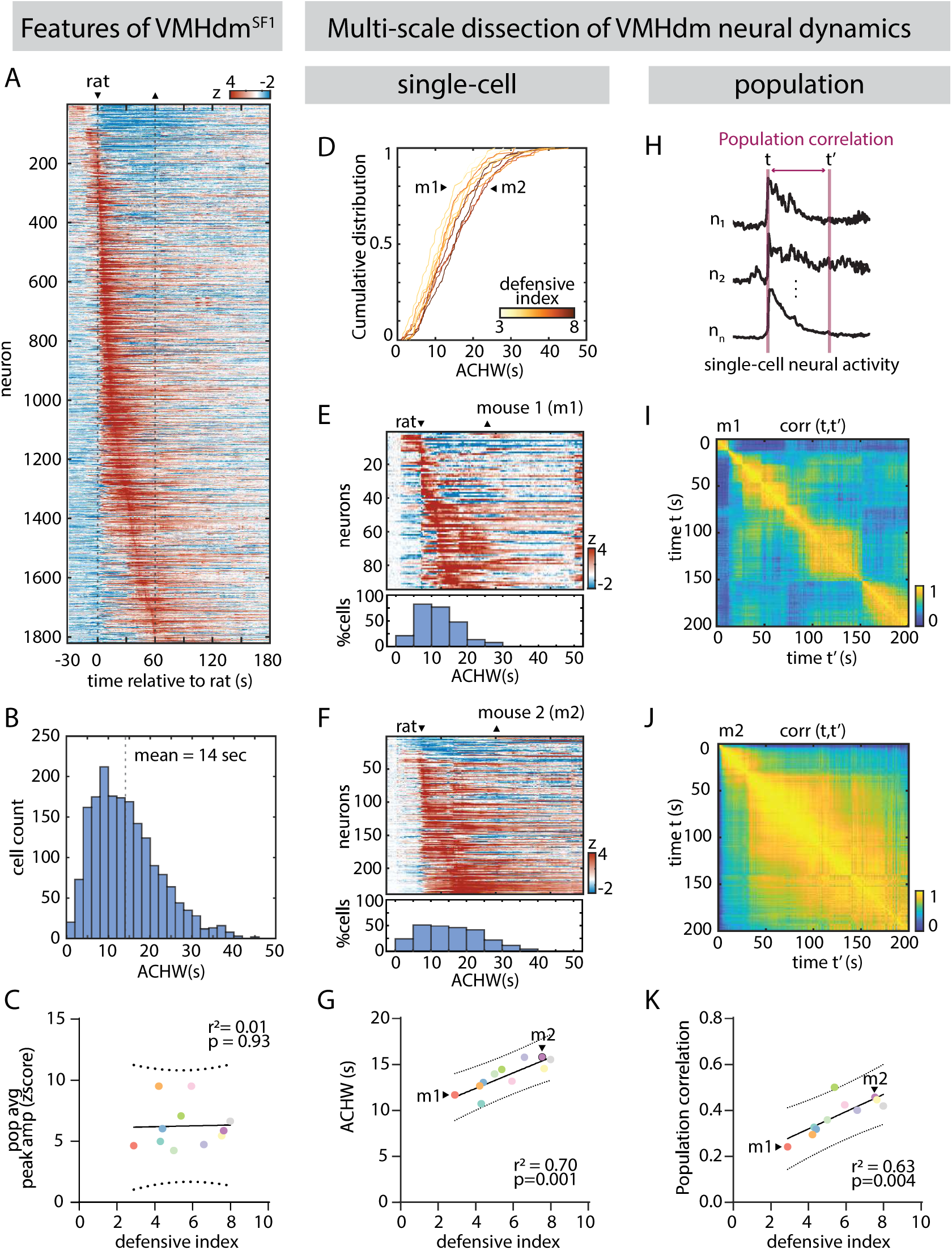
VMHdm dynamics correlate with the individual differences in defensiveness across animals. (A) z-scored neural traces sorted by time of peak from 1821 cells across 11 mice, showing that VMHdm^SF1^ exhibit sequential activity during predator exposure in the PICA test. (B) ACHW distribution for all cells. (C) Correlation between the peak amplitude of VMHdm^SF1^ population average and defensive index (r^2^ = 0.01, p = 0.93). Defensive index (DI) = number of undirected escapes + [percentage of time spent freezing÷10]. (e.g., 3 undirected escapes & 50% time freezing equates to 3 + 5 = DI value of 8). Dashed line indicates 95% confidence interval. (D) Cumulative ACHW distribution of all cells in each individual animal. The color of the line indicates the DI of the respective animal. (E-F) Representative z-scored neural traces indicating time of rat introduction and removal (upper) and corresponding ACHW distribution (lower), from two example animals with (E) low and (F) high DI. (G) Relationship between mean ACHW and the DI for individual mice. (r^2^ = 0.71, p<0.001). Dashed line indicates 95% confidence interval. (H) Diagram illustrating the concept of population correlation. (I-J) Representative population correlation matrices for two animals, one with (I) low and the other with (J) high DI. (K) Relationship between population correlation and DI for individual mice. (r^2^ = 0.63, p<0.01). Dashed line indicates 95% confidence interval. See also Figure S6.

Prior single-cell imaging experiments conducted with head-fixed subjects have shown that individual VMHdm^SF1^ neurons display heterogeneous but reproducible dynamics, resulting in persistent neural activity at the population level which endures beyond the cessation of a threatening stimulus^10^. At the single-cell level, these dynamics were reflected in both sequential and persistent firing, an observation validated in our PICA assays (Figures 6A-B). Intriguingly, however, while all subjects exhibited persistent post-predator VMHdm^SF1^ activity to some degree, there was notable variation in the duration of this persistence (i.e., in its decay kinetics). The observed variance in VMHdm^SF1^ dynamics therefore prompted us to investigate the correlation between VMHdm^SF1^ persistence and individual differences in defensive behaviors.

We correlated the DI of each mouse with different measures of VMHdm^SF1^ dynamics. First, we investigated the relationship between defensive behavior and VMHdm^SF1^ single neurons’ persistent firing, measured as the ACHW (Figures 6B, S1D). This comparison revealed a robust correlation between DI and the mean ACHW across individual animals (r^2^ = 0.70, p = 0.001, n = 11 animals, 165±67 cells/animal; Figures 6D-G). Subjects with high ACHW values exhibited a higher DI index, i.e. more defensive behaviors and vice versa.

Previous studies suggested that in addition to slowly decaying unit firing, collective population mechanisms also play a role in sustaining the overall persistence observed in VMHdm^SF1^ neural dynamics, possibly through mechanisms involving sequential activity (Figure 6A)^10^. Employing a methodology derived from that work^10^, we used population correlation matrices as a surrogate for population-level persistence (Figure 6H). This method presents an alternative way to characterize VMHdm^SF1^ neural dynamics across animals, one that does not rely on the ACHW of individual cells but rather on the correlation between neurons. We again observed a strong correlation between the magnitude of the time-varying population correlation coefficient and DI across individual animals (r^2^ = 0.66, p = 0.004; Figures 6I-K).

Because these correlations were found in data from the PICA assays, we investigated whether the imminence-coding population we identified therein (Figure 5L) also exhibited a similar correlation with DI. To do this, we computed the ACHWs and DIs from the dataset collected using double-reversed PICA experiments (Figure 5E). Consistent with the result obtained using data from animals’ first rat exposure, we again observed a robust correlation (r^2^ = 0.6, p=0.04, n = 7 animals, 192±60 cells/animal) between the mean ACHW computed from all neurons and DI across animals in the double-reversed PICA assays (Figure S6A). Since the population activity of the imminence-coding cells could predict context N vs. F (Figure 5K) and was suggested to encode the level of predator imminence, we expected that this VMHdm^SF1^ subpopulation might exhibit an even stronger correlation with DI. Indeed, the ACHW of the imminence-coding cells exhibited an even higher correlation with the DI (r^2^ = 0.8, p=0.01; Figure S6B), than did the total population (r^2^ = 0.6, p=0.04; Figure S6A). Furthermore, this effect was much weaker in the residual population where no significant correlation was observed (r^2^ = 0.4, p=0.13; Figure S6C).

Although we did not observe any correlation between the peak amplitude of the VMHdm^SF1^ population average and the DI (Figure 6C), we investigated whether the amplitude of the imminence-coding population is correlated with the DI of individual animals. To our surprise, we found that the peak amplitude of the imminence-coding population (normalized to total activity) correlated with the DI across individual animals (r^2^=0.63, p=0.03, Figure S6E), but not that of the total population average (r^2^=0.16, p=0.37; Figure S6D) or the residual population (r^2^=0.12, p=0.43; Figure S6F). This finding further supports the interpretation, based on within-animal differences in activity across (mode N vs. mode F) trials, that the imminence-coding population represents an affective state variable related to predator imminence.

Our unsupervised analysis of neural data revealed that VMHdm^SF1^ neurons encode a broad spectrum of predator-related variables, including threat object identity, a scalable defensive internal state, predator imminence, safety, and novelty or arousal. Using different behavioral assays designed to isolate these variables, we found heterogeneity in their neural representations within the VMHdm^SF1^ population. Determining the extent to which these variables are encoded by overlapping or distinct VMHdm^SF1^ neurons across all assays is challenging, as sequential recordings of the same cell in a single animal subjected to different assays was not feasible due to rapid behavioral habituation of the mice to the predator. To overcome this limitation, we used unsupervised k-means clustering of neural data (performed without any behavioral annotation) acquired during rat presentation to compare response profiles across the different behavioral assays. Clustering performed independently on datasets from the two-chamber, two-chamber with shelter, and PICA assays consistently yielded three physiologically distinct clusters of VMHdm^SF1^ neurons with different response profiles to the rat: cluster 1 cells were persistently activated by rat; cluster 2 cells were transiently activated by rat; and cluster 3 cells were inhibited by rat. Each of these three clusters exhibited a high and specific temporal correlation in its average activity with its cognate cluster across different assays (Figures S7A-C), suggesting that they contain similar populations of cells.

## Discussion

Encounters with a predator require that the prey animal’s brain compute and represent multiple variables and use that information to guide its behavior to maximize its chances for survival. These include threat identification; computation of distance from the predator; mapping possible routes of escape or sites for concealment; generating an internal motive state of fear; and initiating motor programs for specific defensive behaviors. We have used single neuron-resolution calcium imaging in freely behaving mice to investigate which of these predator encounter features are represented by VMHdm^SF1^ neurons, cells which have been shown by perturbation experiments to be necessary and sufficient for defensive behaviors^6,7,13^. Our results provide evidence that diverse features of a predator encounter are represented by physiologically heterogenous subpopulations of VMHdm^SF1^ neurons.

### VMHdm^SF1^ neurons represent multiple internal states

VMHdm^SF1^ neurons are strongly activated in the presence of various predators or predator odors^7,9–12^. Because these stimuli also evoke an internal state of fear or anxiety, it is difficult in principle to disambiguate whether this activity represents predator stimulus identity, internal state or a combination of both, particularly when measuring bulk calcium activity^10,11^. Our k-means clustering of single-cell responses during predator exposure in the 2-chamber assay revealed a major population (cluster 1) that displayed strong responses to a rat but not a toy, which persisted after predator removal. The stimulus-specificity of this response suggests that this population encodes the identity of a threatening object. Consistent with this interpretation, c-Fos labeling studies have shown that different predators activate spatially distinct subsets of VMHdm neurons^9^.

The evidence that cluster 1 responses may also encode an internal defensive state of fear or anxiety is supported by their persistent activity, a feature of such states^6,10,21^. Furthermore, our earlier studies in head-fixed mice exposed to either a rat or an aversive ultrasound stimulus^51^ revealed that the first two principal components of VMHdm^SF1^ activity reflect stimulus identity, and a variable whose value declines with repeated stimulus exposure, respectively^10^. The location of individual units in PC space was dependent on both variables, suggesting that their activity represents a combination of both stimulus identity and internal state.

The k-means analysis also identified a second population (cluster 2) that exhibited robust activation in response to both the rat and the control (toy rat), but which rapidly returned to baseline even while the stimulus was present. The transient and non-discriminative response of this population suggests it may encode novel objects, neophobia, surprise or arousal. The activity of the bulk VMHdm^SF1^ population evoked by a predator therefore represents a combination of these two distinct response patterns. Consequently, the level of predator-evoked fear cannot be inferred simply from the amplitude of total or bulk VMHdm^SF1^ activity.

Prior studies performed using electrophysical recording of anonymous units or calcium imaging of bulk VMHdm^SF1^ population activity in freely behaving animals suggested that the level of activity correlates with predator proximity^11,12^. However, in the two-chamber assay we found that VMHdm^SF1^ neuronal activity was correlated with predator proximity only when the mouse was in the approach phase; this distance-dependent relationship was not maintained following escape. Thus, while VMHdm^SF1^ activity may represent predator proximity during approach, it is not an absolute function of distance to the threat. Instead, our results indicated a similar rate of change in population VMHdm^SF1^ activity over time during both the approach and escape phases, albeit in opposite directions. This observation is consistent with the encoding of an internal state that ramps up during approach and decays following escape with similar kinetics, perhaps reflecting a leaky neural integration function^52^.

Together, our data provide evidence that, in addition to representing threat stimulus identity, VMHdm^SF1^ neurons represent at least two types of internal state evoked by a predator: a threat-specific state of fear or anxiety; and a more general neophobic or arousal response. The mechanisms that regulate the relative magnitudes and stability of these states will be an interesting topic for future study.

### Bi-directional encoding of safety and threat

VMHdm is well-known to be activated by predator cues^7,9–12^. Our unsupervised k-means clustering identified a third subset of VMHdm^SF1^ neurons (cluster 3), whose activity was strongly and persistently suppressed by the predator. Similar cells were identified in previous studies^10,12^. However, those studies did not identify what, if anything, activates those cells. Unexpectedly, we found that these rat-inhibited neurons became activated upon entry into a refugium (shelter), suggesting they encode safety-related information. Thus, VMHdm^SF1^ activity during a predator encounter encodes not only stimulus identity, predator fear and neophobia/arousal but also whether or not the animal is in a safe space, through the activity of physiologically distinct subpopulations.

Safety is typically viewed as antagonistic to threat. Therefore, in principle it could be encoded simply by virtue of reduced activity among threat-encoding cells. Our findings reveal that this is not the case. Instead, we identified distinct subpopulations of VMHdm^SF1^ neurons that represent safety and threat bi-directionally, in an anti-correlated manner. This dichotomous coding could afford animals with fine-scale control over the intensity and valence of their internal state. Since VMHdm also serves as an important metabolic homeostatic center^53,54^, our discovery might shed light on how appetitive behaviors driven by homeostatic needs and defensive responses are balanced in the hypothalamus^55,56^.

How the relative activity of these distinct threat and safety-encoding subpopulations is controlled is not clear. While the data are suggestive of reciprocal inhibitory interactions between them, virtually all VMHdm^SF1^ neurons are glutamatergic^20^. Therefore, if such circuit-level opponency exists, it is likely to be indirect and mediated by inhibitory neurons in other structures, such as the VMH shell, tuberal nucleus (TU), dorsal medial hypothalamus (DMH) or AHN. Studies of conditioned fear have revealed distinct and mutually antagonistic clusters of intercalated inhibitory neurons (ITCs) in the amygdala that encode high- vs. low-fear states^57,58^. Other studies have identified distinct subsets of “fear”- vs. “safety”-encoding cells in the basal amygdala^59^. However, unlike the VMHdm^SF1^ cells, “safety” neurons in the basal amygdala were not inhibited by fear cues, but only responded to the combination of fear + safety cues^59^.

Taken together, our results reveal that even within a genetically defined and anatomically restricted subclass of hypothalamic neurons, cells active during naturalistic predator encounters are functionally diverse and encode distinct information or features relevant to the encounter. Consistent with this finding, single-cell RNA sequencing has revealed that VMHdm^SF1^ neurons contain multiple transcriptomically distinct cell clusters^60,61^. Future research could explore whether each of the functionally distinct VMHdm^SF1^ subpopulations identified here corresponds to a different transcriptomic subtype. If so, it could potentially provide genetic access to these subtypes to investigate their connectivity, physiological properties and causal roles in behavior.

### Population coding of predator imminence

Predator defense behaviors are diverse and include freezing, flight, jumping, risk assessment or undirected “panic-like” escape. These behaviors have been shown to vary according to the distance from the predator and the availability of an escape route^31,32,45^. Such observations have been encapsulated in “Predator Imminence Theory,” which posits that animals adopt distinct defensive strategies as a predator approaches and/or the context changes^28^. How these different behaviors are selected as a function of predator imminence has not been clear. Optogenetic stimulation of VMHdm^SF1^ neurons drives different defensive behaviors in a scalable, threshold-dependent manner^6,13^, but the mechanisms by which it controls this behavioral spectrum remain poorly understood.

The simplest hypothesis is that more intense defensive behaviors would correlate with higher levels of neural activity in a common population, which may ramp up with predator approach. Alternatively, different defensive behaviors could be encoded by distinct and non-overlapping VMHdm^SF1^ subpopulations with different activation thresholds^62^, perhaps with distinct downstream projections^13,63^. Studies of conditioned freezing and flight have shown that these behavioral responses are mediated by distinct and mutually inhibitory neurons in the central amygdala (CEA)^50^. Answering this question for innate defensive responses has been challenging because eliciting freezing by predator exposure in inbred strains of laboratory mice is notoriously difficult^51^. Our novel PICA assay preferentially elicited freezing or undirected escape to the rat, depending on the degree of predator imminence and context^28,31,32,45^. Using this assay, we were able to disambiguate neural representations of sensory cues, motor behavior and internal state by VMHdm^SF1^ neurons.

Strikingly, we found that virtually no VMHdm^SF1^ neurons were tuned specifically to either freezing or undirected escape. In addition, the dominant neural response was independent of the two modes and was explained by the presence of the rat. Efforts to identify subpopulations of neurons tuned to either mode N (near) or mode F (far) context by GLM yielded only a tiny proportion (1%) of such cells. However, by training supervised classifiers on neural activity we were able to identify a subpopulation of neurons whose activity could predict whether the animal was in the N vs. F context. This population comprised ∼34% of VMHdm^SF1^ neurons, indicating that predator imminence is represented in VMHdm by a population code. Notably, the average activity in this subpopulation was higher in the N than in the F context, consistent with the idea that it encodes an internal state that escalates as the probability of a deadly encounter with a predator increases^28^. This suggests that in addition to the three states described earlier— predator induced fear/anxiety, neophobia/arousal, and safety states — a population code represents another distinct state that may be related to the degree of predator imminence. Whether the cells that contribute to this code are related transcriptomically or connectionally is an interesting question for future studies.

Our discovery of the imminence-coding population supports the idea that VMHdm^SF1^ neurons play a key role in representing threat-derived sensory cues and transforming that representation into an internal affective state(s), whose magnitude may control decisions between competing defensive responses. This hypothesis aligns with the anatomical positioning of VMHdm between sensory inputs and motor outputs^3^. VMHdm^SF1^ neurons project to numerous targets, including the anterior hypothalamic nucleus (AHN), paraventricular nucleus (PVN), medial preoptic nucleus (MPN), bed nucleus of stria terminalis (BNST), paraventricular thalamic nucleus (PVT), CEA, medial amygdala (MEA) and PAG^6,13,18^. Identifying the site(s) at which the transformation of state intensity into different types of motor behaviors occurs, and the circuit implementation of this transformation, is a challenging and important future objective.

### Neural correlates of individual variation in defensiveness

We imaged 1800 cells from 11 animals during their first exposure to a rat in their lifetime using the PICA assay and observed wide variation among individual C57BL6/N mice in the quality and quantity of predator-evoked defensive behavioral responses. Intuitively, one might have expected this variation to correlate simply with the overall level of population VMHdm^SF1^ activity. However, that turned out not to be the case: such a correlation was observed, but only among the subset of neurons that contributed to the imminence-coding population. This observation reinforces the conclusion that variable activity within the imminence-coding population represents the level of an internal affective fear-like state, not only within but also across animals.

We also found that not only the magnitude, but also the duration of imminence-coding population’s responses was correlated with the level of defensiveness across animals. However, this feature was a general property of the VMHdm^SF1^ population, although the imminence-coding population was more highly correlated than the total population. Thus, the length of the defensive state appears to be represented by the dynamics of all VMHdm^SF1^ neurons, while the strength of the state is specific to the imminence-coding subpopulation. Previously, we demonstrated a requirement for persistent VMHdm^SF1^ activity in maintaining a persistent state of anxiety^10^. Our current results extend this observation by showing that the duration and amplitude of VMHdm^SF1^ activity scales with the intensity of an internal fear or anxiety state(s). How this persistent neural activity is implemented, scaled, and read out by downstream targets are important questions for the future^10^.

Finally, our finding that the amplitude and duration of activity in the imminence-coding population is correlated with individual variation in defensiveness may be relevant to human psychiatric disorders. Conditions such as post-traumatic stress disorder (PTSD) are often characterized by abnormally strong and long-lasting response towards stimuli^64^. Electrical stimulation of human VMH evokes an emotional “panic attack-like” feeling^65^, suggesting that the role of VMHdm in encoding fear is evolutionarily conserved across species^61,66^. The discovery that a subset of VMHdm^SF1^ neurons encodes the strength and length of fear responses identify them as potential cellular targets for new therapeutic approaches to such disorders.

## Acknowledgements

We thank Y. Huang for genotyping, G. Mancuso and L. Chavarria for administrative assistance, C. Chiu for lab management, K. Lee and the rest of Caltech OLAR staff for animal care and mouse colony management, and members of the Anderson Laboratory for helpful comments on this project. D.J.A. and M.G.S. are Investigators of the Howard Hughes Medical Institute. This work was supported by NIH grants R01MH112593, R01MH123612, and R01NS123916 and by a grant from the Simons Collaboration on the Global Brain to D.J.A. A.N. is supported by the National Science Scholarship from the Agency of Science, Technology and Research, Singapore. K.Y.M.C. is supported by the Croucher Scholarship for Doctoral Study from the Croucher Foundation, Hong Kong.

## Author contributions

K.Y.M.C., A.N., L.L., M.G.S. and D.J.A. contributed to the study design and conceptualization. K.Y.M.C conducted all the experiments and figure preparation. K.Y.M.C. and A.N. performed data analysis. D.J.A. supervised the project. K.Y.M.C. and D.J.A. wrote the manuscript.

## Declaration of interests

The authors declare no competing interests. D.J.A. is a member of the *Neuron* editorial board.

## STAR Methods

### Experimental model and subject details

#### Mice

All experimental procedures involving the use of live mice were carried out in accordance with NIH guidelines and approved by the Institute Animal Care and Use Committee (IACUC) and the Institute Biosafety Committee (IBC) at the California Institute of Technology (Caltech). SF1-Cre mice were obtained from Dr. Brad Lowell ^53^ and maintained as heterozygotes in the Caltech animal facility as described previously; the SF1-Cre line is also available from the Jackson Laboratory (Stock No: 012462). Heterozygous male mice, aged between 8-20 weeks, were used in this study. Since sexual dimorphism has been reported in hypothalamic nuclei such as VMH, we chose to perform all experiments in male mice to eliminate potential effect from the oestrous cycle and maintained consistency with our previous characterization of the VMHdm^SF1^ neurons ^67–69^. All mice were housed in ventilated micro-isolator cages in a temperature-controlled environment (median temperature 23 °C, humidity 60%), under a reversed 11-hour-dark-13-hour light cycle, with ad libitum access to food and water. Mouse cages were changed weekly on a fixed day in which behavioral experiments were not performed. Long-Evans male rates (for use as predators) were obtained from Charles River at 2-3 months of age and were raised to 5-10 months in the Caltech animal facilities.

#### Viruses

SF1-Cre+/- male mice were injected in VMHdm with 100 nl AAV9-syn-FLEX-jGCaMP7f-WPRE (Addgene 104492-AAV9; 2.7 x 10^12^ genomic copies per ml) for microendoscopic calcium imaging.

### Method

#### Behavior tests

All behavioral experiments were performed either in the custom-designed two-chamber arena (Figures 1-3) or the animals’ home cage (Figures 4-6) under red light during the animals’ subjective night phase. Top and front views for the two-chamber arena and front and side views for the home cage were filmed at 30Hz using video recording software, Spinnaker SDK (Teledyne Flir). The position of the animal (center-of-mass) was extracted using EthoVision XT and behaviors were manually scored (Bento^70^).

#### Two-chamber assay

Prior to the introduction of the test animal, the two-chamber arena was lined with fresh bedding. In the two-chamber with shelter assay, an opaque triangular-shaped shelter was placed in the middle of the exploration chamber against the wall. The shelter had a slit on top to allow the Miniscope cable to go through. The tested mouse was introduced to the exploration chamber and allowed to acclimate for 10 minutes followed by 5 minutes of baseline recording. Then a toy rat of a similar size to the Long-Evans male was introduced into the stimulus chamber. The toy rat remained there for 5 minutes then it was removed. After another 5 minutes, a live rat was introduced into the stimulus chamber for 5 minutes.

#### Predator threat context assay (PICA)

Prior to the test day, the animals were acclimated in the test room in their home cage for one hour every day for four days. Cage change occurred at least 5 days before the test day. During both acclimation and behavioral experiments, the cage lid was removed, and a rectangular-shaped acrylic frame was placed above the cage. The frame was used to extend the height of the cage wall so that the mice were not able to jump out of the cage during predator exposure. On the test day, the mice were allowed to acclimate in this open-lid configuration for 30 minutes prior to rat introduction to reduce baseline anxiety behaviors, such as thigmotaxis and bedding digging. Then a live rat in a plastic enclosure was lowered into the mouse’s home cage by the experimenter. During transition between mode N and mode F, the rat enclosure was manually moved by the experimenter.

#### Stereotaxic surgery

Surgeries were performed on adult SF1-Cre+/- mice aged 8-10 weeks. Mice were anesthetized with 5% isoflurane and placed on a stereotaxic apparatus (David Kopf Instruments). The virus was injected into VMHdm using a pulled glass capillary (World Precision Instruments) and a pressure injector (Micro4 controller, World Precision Instruments), at a flow rate of 40nL/min. The injection volume was 100nL. The stereotaxic injection coordinates were based on a high resolution three-dimensional surgical atlas^71^ (VMHdm, anterior-posterior: -4.6, medial-lateral: ±0.4, dorsal-ventral: -5.6). The GRIN lens used for microendoscopic imaging was slowly lowered into the brain to 100μm above the virus injection site and fixed on the skull with dental cement (Parkell). After surgery, mice were allowed to recover for at least 6 weeks before behavioral testing.

#### Microendoscopic imaging data acquisition

Single-cell calcium imaging data was acquired at 30Hz with the following parameters: 2x spatial down sampling, 0.2-0.4 light-emitting diode power, and 1-5x gain. The parameters were calibrated 3 days before behavioral testing depending on the GCaMP expression level indicated by the image histogram in Inscopix Data Acquisition Software. The data acquisition box (DAQ, Inscopix) emitted a transistor-transistor logic (TTL) pulse to synchronize behavioral video frame acquisition by Spinnaker SDK (Teledyne Flir).

#### Histology

Upon finishing all behavioral experiments, virus expression and GRIN lens placement were determined histologically. Mice were transcranial perfused with 1x PBS at room temperature, followed by 4% paraformaldehyde (PFA). Brains were harvested and post-fixed in 4% PFA for 24 hours at 4°C and subsequently transferred to 30% sucrose/PBS at 4°C for another 48 hours. Brains were then embedded in OCT mounting medium and frozen at -80 °C for sectioning using a cryostat (Leica Biosystems). Brain slices (thickness: 60μm) were obtained and stained with DAPI/PBS (0.5μg/ml) for 15 minutes at room temperature. The counterstained brain slides were then mounted on Superfrost slides and imaged with an epifluorescent microscope (Olympus VS120).

#### Quantification and statistical analysis

Statistical analysis was performed with GraphPad PRISM (GraphPad Software). Sample distribution normality was first determined with Kolmogorov-Smirnov tests. The Wilcoxon rank test was used for experiments with paired samples and the Mann-Whitney U-test for unpaired samples.

#### Microendoscopic data preprocessing and unit extraction

Motion correction, 2x spatial down-sampling and 3x temporal down-sampling to 10Hz were performed using Inscopix Data Processing Software. After preprocessing, calcium traces were extracted and deconvolved using the CNMF-E large data pipeline with the following parameters: patch_dims = [32, 32], gSig = 3, gSiz = 13, ring_radius = 19, min_corr = 0.7, min_pnr = 8 ^72^. Manual iterations were performed on the CNMF-E output to remove outliers. The extracted neural traces were z-scored before analysis.

#### Choice probability

Choice probability is used to determine whether a single neuron might be probabilistically tuned to either of two behaviors. This calculation was performed as described in previous studies ^36^. In cases where only one behavior was observed, we subtracted the time frames in which the behavior occurred from the period of rat presence to use as the comparative behavior. Choice probability values range from 0 to 1, where a value of 0.5 indicates that the neuron’s activity does not distinguish between the two behaviors. To be considered tuned to a particular behavior, a neuron must exhibit a choice probability of less than 0.3 or greater than 0.7 and above 2σ or below -2σ of the choice probability computed using shuffled data.

#### Generalized linear model

To predict neural activity from freezing and undirected escape, we trained generalized linear models as previously described to predict the activity of each neuron as a weight linear combination of the rat presence and freezing or undirected escape ^40^. All behaviors were provided as binarized vectors and convolved using a behavior-filter that described how a neuron integrates stimuli over a 10-second period. The model was fitted using 10-fold cross-validation, and model performance was reported as cross-validated R^2^.

#### Decoder analysis

We constructed frame-wise linear SVM decoders (as described previously) to discriminate mode A and mode B from neural data ^42,43^. Briefly, manual annotations of each mode were used to provide training labels to an SVM decoder. We generated bar graphs depicting decoder accuracy, which discriminated between the two modes based on the imaged activity in individual frames (sampled at 10 Hz). Equal numbers of mode A and mode B frames (frame-wise decoder) were used during decoder training, to ensure a chance decoder performance of 50%. ‘Shuffled’ decoder data were generated by training the decoder on the same neural data, but with mode annotations randomly assigned to each bout.

Decoding was repeated 20 times for each imaged mouse, and decoder performance was reported as the average accuracy across imaged mice. For significance testing, the mean accuracy of the decoder trained on shuffled data was computed across mice, in each condition, and shuffling was repeated 1,000 times. Significance was determined across imaged mice using the Mann–Whitney U test between the mean accuracy of the decoders trained on real versus shuffled data.

#### Autocorrelation half-width

We computed autocorrelation halfwidths by calculating the autocorrelation function for each neuron’s time-series data (y^t^) for a set of lags as described previously ^36^. Briefly, for a time series (y^t^), the autocorrelation for lag k is:

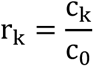

Where c_k_ is defined as:

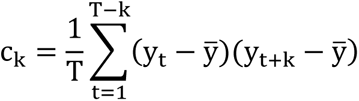

where ȳ is the average value of the time series across the interval from t to t + k, c_0_ is the sample variance of the data, and T is the number of timepoints. The half-width is found for each neuron as the point where the autocorrelation function reaches a value of 0.5 (Figure S1D).

#### Population correlation

To quantify the correlation across neurons at a population level, we computed the autocorrelation matrix of population activity between all pairs of time points, t and t′, in a 100-s interval following rat presentation as previously performed^10^. We then quantified time-evolving dynamics by taking the mean autocorrelation as a function of the lag between time points |t – t′|. To compare this metric across mice, we used the mean autocorrelation at a lag of 50 seconds.

## Supplemental Figure Legends

**Figure S1. Additional characterization of the two-chamber assay, related to Figures 1 and 2**

(A-C) (A) Peak amplitude, (B) area under the curve, and (C) decay constant of VMHdm^SF1^ population average during rat and toy presentation

(D) Autocorrelation function of 3 example cells with (top) high, (middle) medium, and (bottom) low ACHW values.

(E) Venn diagram showing the overlap of rat and toy-activated cells, where “activation” is defined as a response >2σ above baseline. Rat-activated: 74.4%, toy-activated:57.3%, overlap: 43.7%.

(F) Choice probability histogram and percentages of rat-tuned (27%; red) and toy-tuned (16%; green) cells. The mixed population contains cells that are activated by both or neither of the stimuli. (G-H) Mean ± s.e.m. population responses of rat-tuned and toy-tuned cells aligned to (G) toy and

(H) rat introductions.

(I-J) Comparison of VMHdm^SF1^ population average activity level between zone I and zone VI (see main Figure 2F) during (I) approach and (J) escape. \*\*\*\**P*<0.0001, ns: not significant.

(K) Mean ± s.e.m. response of individual VMHdm^SF1^ neurons during rat investigation (black line; duplicated from Figure 2B for comparative purposes), overlaid with velocity of the animal (green).

(L) Fraction of variance in VMHdm^SF1^ neural activity explained by distance to rat, speed, presence in inves or distal zones.

**Figure S2. Additional characterization of the two-chamber assay with shelter, related to Figure 3**

(A-B) Example traces of 5 (A) shelterIN and (B) shelterOUT cells aligned to shelter entry.

(C) Cell raster and k-means clustering of neural dynamics during rat and toy presentation in the two-chamber with shelter assay (duplicated from Figure 3L for comparative purposes). Red: cluster 1; Blue: cluster 2; Brown: cluster 3.

(D) Mean ± s.e.m. population responses of VMHdm^SF1^ single cells in each cluster during rat and toy presentations.

(E-G) (E) Peak/trough amplitude, (F) average activity, and (G) ACHW of VMHdm^SF1^ single cells in each cluster during rat and toy presentations. ns: not significant, \*\*\**P*<0.001, \*\*\*\**P*<0.0001.

**Figure S3. Additional characterization of PICA, related to Figures 3 and 4**

(A) Schematics of behavior paradigm in wild-type non-surgical (WTNS) animals.

(B) % time spent freezing in mode N and F by WTNS animals (n = 7 mice).

(C) Number of jumps displayed in mode N and F by WTNS animals.

(D-E) Percentage of time spent freezing in (D) mode N and (E) mode F by WTNS and Miniscope-implanted SF1-Cre mice. ns: not significant. (WTNS: n = 7 mice; Miniscope: n = 11 mice)

(F) Schematics and presentation sequence of the PICA test (duplicated from Figure 4C for comparative purposes).

(G) Percentage of time spent freezing in mode N and F during N>F (left) and F>N (right) respectively.

(H) Number of undirected escapes displayed in mode N and F during N>F (left) and F>N (right) respectively.

Representative z-scored single-cell neural traces and population average with behavioral annotations during PICA test. Green shaded area: freezing; orange shaded area: undirected escape. (J-K) Mean population responses of VMHdm^SF1^ cells aligned to the onset of (J) freezing and (K) undirected escape. (n = 11 mice)

(L) Mean activity of VMHdm^SF1^ neurons during 2 seconds before the onset of behaviors (pre), during the behavior (beh) and 2 seconds after the offset of behaviors (post) for freezing (left) and undirected escape (right).

(M) The fraction of variance in VMHdm^SF1^ single-cell neural activity explained by rat presentation (mode N + mode F) was first calculated and then removed from total neural activity by regression. Next, the fraction of variance in the residual neural activity explained by freezing and undirected escape was calculated. (n = 1821 cells, 11 mice)

(N) GLM architecture for detecting cells that were selectively tuned to freezing or undirected escape. A separate GLM was implemented for each behavior, training each neuron with behavior annotations for freezing/undirected escape and rat presence (Mode N and Mode F) as regressors, along with behavior filters. A non-negativity constraint was applied to aid the interpretation of GLM weights.

(O) Fraction of cells selectively tuned to rat presence and behaviors (upper: undirected escape; lower: freezing), as identified by GLM analysis.

**Figure S4. Both PICA modes induce indistinguishable bulk-level activity and dynamics, related to Figure 5**

(A) Schematics of the behavior paradigm in which each animal was presented with either mode N or F for 60 seconds, repeated three times at 10-minute intervals. Separate groups of animals were used for each test to avoid habituation.

(B) Mean ± s.e.m. population responses of VMHdm^SF1^ neurons during mode N (red) and F (blue) averaged across three bouts.

(C) Mean ± s.e.m. population response of individual VMHdm^SF1^ neurons during mode N and F for each bout.

(D) Peak amplitude of VMHdm^SF1^ population average in response to rat introduction (first peak) average across three bouts for mode N and F. Mean ± s.e.m.

(E) Latency to first peak for VMHdm^SF1^ population average averaged across three bouts for mode N and F. Mean ± s.e.m.

(F) Latency to first peak of VMHdm^SF1^ population average across three bouts for mode N and F. Mean ± s.e.m.

(G) The difference in latency to first peak between the first bout and third bout in mode N and B.

(H) Fraction of cells that are activated (defined by>2σ above baseline:), inhibited (defined by <2σ below baseline:) or non-responsive (-2σ<x<2σ) by rat in both mode N and F.

(I) Mean percentage of activated (left, violet shading) and inhibited (right, orange shading) cells averaged across three bouts in mode N and F.

(J) Percentage of activated and inhibited cell across three bouts in mode N and F. Mean ± s.e.m.; solid line: activated population; dashed line: inhibited population.

(K) Mean ± s.e.m. responses of activated and inhibited population populations averaged across three bouts for mode N and mode F.

(L) Mean peak amplitude of activated (left) and mean trough amplitude of inhibited (right) populations averaged across three bouts for mode N and F.

(M) Mean ± s.e.m. responses of activated and inhibited populations across three bouts for mode N and F.

(N) Peak amplitude of activated population and trough amplitude of inhibited population across three bouts for mode N and F. Mean ± s.e.m.

(O) Schematics and presentation sequence of the PICA (duplicated from Figure 4C for comparative purposes).

(P) Peak amplitude of VMHdm^SF1^ population average activity for each mode during N>F (left) and F>N (right). ns: not significant; \*\**P*<0.01.

(Q) Percentage of activated cells for each mode during (left) N>F and (right) F>N. \*\**P*<0.01.

(R) Decay constant of VMHdm^SF1^ bulk activity during N>F and F>N. ns: not significant.

**Figure S5. Additional characterization of the double-reversed PICA and imminence-coding population, related to Figure 5**

(A) Representative z-scored neural traces of imminence-coding cells from an example animal.

(B) Average activity and (C) peak amplitude of imminence-coding cells during each mode. (n = 7 mice)

Left: Fraction of cells weighted by the modes N vs. F decoder (duplicated from Figure 6K (left) for comparative purposes). Right: Mean ± s.e.m. population responses of residual cells (i.e., non-decoder weighted) during PICA.

(E) Average activity and (F) peak amplitude of mean activity of residual cells averaged within each animal. ns: not significant, \**P*<0.05.

(G) Average activity and (H) peak amplitude of single residual cells. \*\*\*\**P*<0.0001.

(I) Mean responses of the imminence-coding cells at each given distance (binned every 4 cm) during mode F.

(J) Upper: Mean ± s.e.m. responses of VMHdm^SF1^ single cells in each cluster (solid lines) during double-reversed PICA overlaid with the imminence-coding population (black dashed line, duplicated from Figure 5L for comparative purposes). Lower: Cell raster with k-means clustering of neural dynamics during PICA. Red: cluster 1; Blue: cluster 2; Brown: cluster 3.

(K) Imminence-coding population composition by k-means clusters.

(L) Non-negative linear decoder (SVM) architecture for predicting mode N versus mode F, trained on VMHdm^SF1^ neural activity from cluster 1 only.

(M) Performance of SVM decoder in differentiating defensive modes trained on cluster 1 cells. \*\*\**P*<0.001.

(N) Mean ± s.e.m. responses of cluster 1 cells weighted by the decoder during double-reversed PICA.

**Figure S6. Additional characterization of VMHdmSF1 neural dynamics and its relationship with DI, analyzed using data only from double-reversed PICA assay, related to Figures 5 and 6**

(A-C) Correlation between the mean ACHW of (A) all (double-reversed PICA) cells (r^2^ = 0.6, p < 0.05), (B) imminence-coding cells (r^2^ = 0.8, p < 0.01), (C) residual cells (r^2^ = 0.4, p = 0.13) and DI (n = 7 mice). Dashed line indicates 95% confidence interval.

(D-F) Correlation between the peak amplitude of the population average of (D) all cells (r^2^ = 0.16, p = 0.37), (E) imminence-coding cells (r^2^ = 0.63, p < 0.1), (F) residual cells (r^2^ = 0.12, p = 0.43) and DI. Dashed line indicates 95% confidence interval.

**Figure S7. Temporal correlation between k-means clusters across different assays, related to Figures 1, 3, and 5**

(A-C) Correlation (r value) between k-means clusters 1 – 3 identified in (A) two-chamber assays and two-chamber with shelter assays, (B) two-chamber assays and PICA assays, and (C) two-chamber with shelter assays and PICA assays.

## Notes

### Competing Interest Statement

The authors have declared no competing interest.

## References

1. Rosen, J.B. (2004). The neurobiology of conditioned and unconditioned fear: a neurobehavioral system analysis of the amygdala. Behav. Cogn. Neurosci. Rev. 3, 23–41.

2. LeDoux, J. (2012). Rethinking the emotional brain. Neuron 73, 1052.

3. Gross, C.T., and Canteras, N.S. (2012). The many paths to fear. Nat. Rev. Neurosci. 13, 651– 658.

4. Adolphs, R. (2013). The biology of fear. Curr. Biol. 23, R79–93.

5. Canteras, N.S. (2002). The medial hypothalamic defensive system: hodological organization and functional implications. Pharmacol. Biochem. Behav. 71, 481–491.

6. Kunwar, P.S., Zelikowsky, M., Remedios, R., Cai, H., Yilmaz, M., Meister, M., and Anderson, D.J. (2015). Ventromedial hypothalamic neurons control a defensive emotion state. Elife 4. 10.7554/eLife.06633.

7. Silva, B.A., Mattucci, C., Krzywkowski, P., Murana, E., Illarionova, A., Grinevich, V., Canteras, N.S., Ragozzino, D., and Gross, C.T. (2013). Independent hypothalamic circuits for social and predator fear. Nat. Neurosci. 16, 1731–1733.

8. Silva, B.A., Gross, C.T., and Gräff, J. (2016). The neural circuits of innate fear: detection, integration, action, and memorization. Learn. Mem. 23, 544–555.

9. Carvalho, V.M. de A., Nakahara, T.S., Souza, M.A. de A., Cardozo, L.M., Trintinalia, G.Z., Pissinato, L.G., Venancio, J.O., Stowers, L., and Papes, F. (2020). Representation of Olfactory Information in Organized Active Neural Ensembles in the Hypothalamus. Cell Rep. 32, 108061.

10. Kennedy, A., Kunwar, P.S., Li, L.-Y., Stagkourakis, S., Wagenaar, D.A., and Anderson, D.J. (2020). Stimulus-specific hypothalamic encoding of a persistent defensive state. Nature 586, 730–734.

11. Tobias, B.C., Schuette, P.J., Maesta-Pereira, S., Torossian, A., Wang, W., Sethi, E., and Adhikari, A. (2023). Characterization of ventromedial hypothalamus activity during exposure to innate and conditioned threats. Eur. J. Neurosci. 10.1111/ejn.15937.

12. Esteban Masferrer, M., Silva, B.A., Nomoto, K., Lima, S.Q., and Gross, C.T. (2020). Differential Encoding of Predator Fear in the Ventromedial Hypothalamus and Periaqueductal Grey. J. Neurosci. 40, 9283–9292.

13. Wang, L., Chen, I.Z., and Lin, D. (2015). Collateral pathways from the ventromedial hypothalamus mediate defensive behaviors. Neuron 85, 1344–1358.

14. Blanchard, D.C., Williams, G., Lee, E.M.C., and Blanchard, R.J. (1981). Taming of wild Rattus norvegicus by lesions of the mesencephalic central gray. Psychobiology.

15. Fuchs, S.A., Edinger, H.M., and Siegel, A. (1985). The organization of the hypothala-mic pathways mediating affective defense behavior in the cat. Brain Res.

16. McNaughton, N. (2011). Fear, anxiety and their disorders: Past, present and future neural theories. Psychology and Neuroscience 4, 173–181.

17. Swanson, L.W. (2000). Cerebral hemisphere regulation of motivated behavior. Brain Res 886, 113–164.

18. Canteras, N.S., Simerly, R.B., and Swanson, L.W. (1994). Organization of projections from the ventromedial nucleus of the hypothalamus: a Phaseolus vulgaris-leucoagglutinin study in the rat. J. Comp. Neurol. 348, 41–79.

19. Canteras, N.S., Simerly, R.B., and Swanson, L.W. (1995). Organization of projections from the medial nucleus of the amygdala. a PHAL study in the rat. J Comp Neurol 360, 213–245.

20. Tong, Q., Ye, C., McCrimmon, R.J., Dhillon, H., Choi, B., Kramer, M.D., Yu, J., Yang, Z., Christiansen, L.M., Lee, C.E., et al. (2007). Synaptic glutamate release by ventromedial hypothalamic neurons is part of the neurocircuitry that prevents hypoglycemia. Cell Metab. 5, 383–393.

21. Anderson, D.J., and Adolphs, R. (2014). A framework for studying emotions across species. Cell 157, 187–200.

22. Campos, C.A., Bowen, A.J., Roman, C.W., and Palmiter, R.D. (2018). Encoding of danger by parabrachial CGRP neurons. Nature 555, 617–622.

23. Kang, S.J., Liu, S., Ye, M., Kim, D.-I., Pao, G.M., Copits, B.A., Roberts, B.Z., Lee, K.-F., Bruchas, M.R., and Han, S. (2022). A central alarm system that gates multi-sensory innate threat cues to the amygdala. Cell Rep. 40, 111222.

24. Grewe, B.F., Gründemann, J., Kitch, L.J., Lecoq, J.A., Parker, J.G., Marshall, J.D., Larkin, M.C., Jercog, P.E., Grenier, F., Li, J.Z., et al. (2017). Neural ensemble dynamics underlying a long-term associative memory. Nature 543, 670–675.

25. Falkner, A.L., Dollar, P., Perona, P., Anderson, D.J., and Lin, D. (2014). Decoding ventromedial hypothalamic neural activity during male mouse aggression. JNeurosci 34, 5971– 5984.

26. Marques, J.C., Li, M., Schaak, D., Robson, D.N., and Li, J.M. (2020). Internal state dynamics shape brainwide activity and foraging behaviour. Nature 577, 239–243.

27. Schaffer, E.S., Mishra, N., Whiteway, M.R., Li, W., Vancura, M.B., Freedman, J., Patel, K.B., Voleti, V., Paninski, L., Hillman, E.M.C., et al. (2023). The spatial and temporal structure of neural activity across the fly brain. Nat. Commun. 14, 5572.

28. Fanselow, M.S., and Lester, L.S. (1988). A functional behavioristic approach to aversively motivated behavior: Predatory imminence as a determinant of the topography of defensive behavior. In Evolution and learning, (pp, R. C. Bolles, ed. (Lawrence Erlbaum Associates, Inc, xi), pp. 185–212.

29. Fanselow, M.S. (1994). Neural organization of the defensive behavior system responsible for fear. Psychon. Bull. Rev. 1, 429–438.

30. Blanchard, D.C., Blanchard, R.J., and Griebel, G. (2005). Defensive responses to predator threat in the rat and mouse. Curr. Protoc. Neurosci. Chapter 8, Unit 8.19.

31. Blanchard, D.C., Griebel, G., and Blanchard, R.J. (2003). The Mouse Defense Test Battery: pharmacological and behavioral assays for anxiety and panic. Eur. J. Pharmacol. 463, 97–116.

32. Blanchard, D.C., Griebel, G., and Blanchard, R.J. (2001). Mouse defensive behaviors: pharmacological and behavioral assays for anxiety and panic. Neurosci. Biobehav. Rev. 25, 205–218.

33. Mobbs, D., Yu, R., Rowe, J.B., Eich, H., FeldmanHall, O., and Dalgleish, T. (2010). Neural activity associated with monitoring the oscillating threat value of a tarantula. Proc. Natl. Acad. Sci. U. S. A. 107, 20582–20586.

34. Mobbs, D., Petrovic, P., Marchant, J.L., Hassabis, D., Weiskopf, N., Seymour, B., Dolan, R.J., and Frith, C.D. (2007). When fear is near: threat imminence elicits prefrontal-periaqueductal gray shifts in humans. Science 317, 1079–1083.

35. Ziv, Y., Burns, L.D., Cocker, E.D., Hamel, E.O., Ghosh, K.K., Kitch, L.J., El Gamal, A., and Schnitzer, M.J. (2013). Long-term dynamics of CA1 hippocampal place codes. Nat. Neurosci. 16, 264–266.

36. Remedios, R., Kennedy, A., Zelikowsky, M., Grewe, B.F., Schnitzer, M.J., and Anderson, D.J. (2017). Social behaviour shapes hypothalamic neural ensemble representations of conspecific sex. Nature 550, 388–392.

37. Ghosh, K.K., Burns, L.D., Cocker, E.D., Nimmerjahn, A., Ziv, Y., Gamal, A.E., and Schnitzer, M.J. (2011). Miniaturized integration of a fluorescence microscope. Nat. Methods 8, 871– 878.

38. Cavanagh, S.E., Towers, J.P., Wallis, J.D., Hunt, L.T., and Kennerley, S.W. (2018). Reconciling persistent and dynamic hypotheses of working memory coding in prefrontal cortex. Nat. Commun. 9, 3498.

39. Murray, J.D., Bernacchia, A., Freedman, D.J., Romo, R., Wallis, J.D., Cai, X., Padoa-Schioppa, C., Pasternak, T., Seo, H., Lee, D., et al. (2014). A hierarchy of intrinsic timescales across primate cortex. Nat. Neurosci. 17, 1661–1663.

40. Liu, M., Nair, A., Linderman, S.W., and Anderson, D.J. (2023). Periodic hypothalamic attractor-like dynamics during the estrus cycle. bioRxivorg. 10.1101/2023.05.22.541741.

41. Lutas, A., Kucukdereli, H., Alturkistani, O., Carty, C., Sugden, A.U., Fernando, K., Diaz, V., Flores-Maldonado, V., and Andermann, M.L. (2019). State-specific gating of salient cues by midbrain dopaminergic input to basal amygdala. Nat. Neurosci. 22, 1820–1833.

42. Karigo, T., Kennedy, A., Yang, B., Liu, M., Tai, D., Wahle, I.A., and Anderson, D.J. (2021). Distinct hypothalamic control of same- and opposite-sex mounting behaviour in mice. Nature 589, 258–263.

43. Yang, B., Karigo, T., and Anderson, D.J. (2022). Transformations of neural representations in a social behaviour network. Nature 608, 741–749.

44. Remedios, R., Kennedy, A., Zelikowsky, M., Grewe, B.F., Schnitzer, M.J., and Anderson, D.J. (2017). Social behaviour shapes hypothalamic neural ensemble representations of conspecific sex. Nature 550, 388–392.

45. Blanchard, R.J., Hebert, M.A., Ferrari, P.F., Palanza, P., Figueira, R., Blanchard, D.C., and Parmigiani, S. (1998). Defensive behaviors in wild and laboratory (Swiss) mice: the mouse defense test battery. Physiol. Behav. 65, 201–209.

46. Blanchard, R.J., Flannelly, K.J., and Blanchard, D.C. (1986). Defensive behaviors of laboratory and wild Rattus norvegicus. J. Comp. Psychol. 100, 101–107.

47. Bang, J.Y., Sunstrum, J.K., Garand, D., Parfitt, G.M., Woodin, M., Inoue, W., and Kim, J. (2022). Hippocampal-hypothalamic circuit controls context-dependent innate defensive responses. Elife 11. 10.7554/eLife.74736.

48. Fanselow, M.S., Hoffman, A.N., and Zhuravka, I. (2019). Timing and the transition between modes in the defensive behavior system. Behav. Processes 166, 103890.

49. Kim, E.J., Park, M., Kong, M.-S., Park, S.G., Cho, J., and Kim, J.J. (2015). Alterations of Hippocampal Place Cells in Foraging Rats Facing a “Predatory” Threat. Curr. Biol. 25, 1362– 1367.

50. Fadok, J.P., Krabbe, S., Markovic, M., Courtin, J., Xu, C., Massi, L., Botta, P., Bylund, K., Müller, C., Kovacevic, A., et al. (2017). A competitive inhibitory circuit for selection of active and passive fear responses. Nature 542, 96–100.

51. Mongeau, R., Miller, G.A., Chiang, E., and Anderson, D.J. (2003). Neural Correlates of Competing Fear Behaviors Evoked by an Innately Aversive Stimulus. J. Neurosci. 23, 3855– 3868.

52. Goldman, M.S., Compte, A., and Wang, X.-J. (2009). Neural Integrator Models. In Encyclopedia of Neuroscience, L. R. Squire, ed. (Academic Press), pp. 165–178.

53. Dhillon, H., Zigman, J.M., Ye, C., Lee, C.E., McGovern, R.A., Tang, V., Kenny, C.D., Christiansen, L.M., White, R.D., Edelstein, E.A., et al. (2006). Leptin directly activates SF1 neurons in the VMH, and this action by leptin is required for normal body-weight homeostasis. Neuron 49, 191–203.

54. Klöckener, T., Hess, S., Belgardt, B.F., Paeger, L., Verhagen, L.A.W., Husch, A., Sohn, J.-W., Hampel, B., Dhillon, H., Zigman, J.M., et al. (2011). High-fat feeding promotes obesity via insulin receptor/PI3K-dependent inhibition of SF-1 VMH neurons. Nat. Neurosci. 14, 911– 918.

55. Canteras, N.S. (2018). Hypothalamic survival circuits related to social and predatory defenses and their interactions with metabolic control, reproductive behaviors and memory systems. Curr. Opin. Behav. Sci. 24, 7–13.

56. Tseng, Y.-T., Schaefke, B., Wei, P., and Wang, L. (2023). Defensive responses: behaviour, the brain and the body. Nat. Rev. Neurosci. 10.1038/s41583-023-00736-3.

57. Hagihara, K.M., Bukalo, O., Zeller, M., Aksoy-Aksel, A., Karalis, N., Limoges, A., Rigg, T., Campbell, T., Mendez, A., Weinholtz, C., et al. (2021). Intercalated amygdala clusters orchestrate a switch in fear state. Nature 594, 403–407.

58. Busti, D., Geracitano, R., Whittle, N., Dalezios, Y., Mańko, M., Kaufmann, W., Sätzler, K., Singewald, N., Capogna, M., and Ferraguti, F. (2011). Different fear states engage distinct networks within the intercalated cell clusters of the amygdala. J. Neurosci. 31, 5131–5144.

59. Sangha, S., Chadick, J.Z., and Janak, P.H. (2013). Safety encoding in the basal amygdala. J. Neurosci. 33, 3744–3751.

60. Kim, D.-W., Yao, Z., Graybuck, L.T., Kim, T.K., Nguyen, T.N., Smith, K.A., Fong, O., Yi, L., Koulena, N., Pierson, N., et al. (2019). Multimodal Analysis of Cell Types in a Hypothalamic Node Controlling Social Behavior. Cell 179, 713–728.e17.

61. Affinati, A.H., Sabatini, P.V., True, C., Tomlinson, A.J., Kirigiti, M., Lindsley, S.R., Li, C., Olson, D.P., Kievit, P., Myers, M.G., et al. (2021). Cross-species analysis defines the conservation of anatomically segregated VMH neuron populations. Elife 10. 10.7554/eLife.69065.

62. Evans, D., Vanessa, A., Vale, R., Ruehle, S., Lefler, Y., and Branco, T. (2018). A synapticthreshold mechanism for computing escape decisions. Nature 558, 590–594.

63. Ma, J., du Hoffmann, J., Kindel, M., Beas, B.S., Chudasama, Y., and Penzo, M.A. (2021). Divergent projections of the paraventricular nucleus of the thalamus mediate the selection of passive and active defensive behaviors. Nat. Neurosci. 10.1038/s41593-021-00912-7.

64. Dymond, S., Dunsmoor, J.E., Vervliet, B., Roche, B., and Hermans, D. (2015). Fear Generalization in Humans: Systematic Review and Implications for Anxiety Disorder Research. Behav. Ther. 46, 561–582.

65. Wilent, W.B., Oh, M.Y., Buetefisch, C.M., Bailes, J.E., Cantella, D., Angle, C., and Whiting, D.M. (2010). Induction of panic attack by stimulation of the ventromedial hypothalamus. J. Neurosurg. 112, 1295–1298.

66. Mobbs, D., Marchant, J.L., Hassabis, D., Seymour, B., Tan, G., Gray, M., Petrovic, P., Dolan, R.J., and Frith, C.D. (2009). From threat to fear: the neural organization of defensive fear systems in humans. J. Neurosci. 29, 12236–12243.

67. Xu, X., Coats, J.K., Yang, C.F., Wang, A., Ahmed, O.M., Alvarado, M., Izumi, T., and Shah, N.M. (2012). Modular genetic control of sexually dimorphic behaviors. Cell 148, 1066–1067.

68. Yang, C.F., Chiang, M.C., Gray, D.C., Prabhakaran, M., Alvarado, M., Juntti, S.A., Unger, E.K., Wells, J.A., and Shah, N.M. (2013). Sexually dimorphic neurons in the ventromedial hypothalamus govern mating in both sexes and aggression in males. Cell 153, 896–909.

69. Correa, S.M., Newstrom, D.W., Warne, J.P., Flandin, P., Cheung, C.C., Lin-Moore, A.T., Pierce, A.A., Xu, A.W., Rubenstein, J.L., and Ingraham, H.A. (2015). An estrogen-responsive module in the ventromedial hypothalamus selectively drives sex-specific activity in females. Cell Rep. 10, 62–74.

70. Segalin, C., Williams, J., Karigo, T., Hui, M., Zelikowsky, M., Sun, J.J., Perona, P., Anderson, D.J., and Kennedy, A. (2021). The Mouse Action Recognition System (MARS) software pipeline for automated analysis of social behaviors in mice. Elife 10. 10.7554/eLife.63720.

71. Chan, E., Kovacevíc, N., Ho, S.K.Y., Henkelman, R.M., and Henderson, J.T. (2007). Development of a high resolution three-dimensional surgical atlas of the murine head for strains 129S1/SvImJ and C57Bl/6J using magnetic resonance imaging and micro-computed tomography. Neuroscience 144, 604–615.

72. Zhou, P., Resendez, S.L., Rodriguez-Romaguera, J., Jimenez, J.C., Neufeld, S.Q., Giovannucci, A., Friedrich, J., Pnevmatikakis, E.A., Stuber, G.D., Hen, R., et al. (2018). Efficient and accurate extraction of in vivo calcium signals from microendoscopic video data. Elife 7. 10.7554/elife.28728.

