## Supplementary figures and images for "Population coding of predator imminence in the hypothalamus"

### supplementary figures

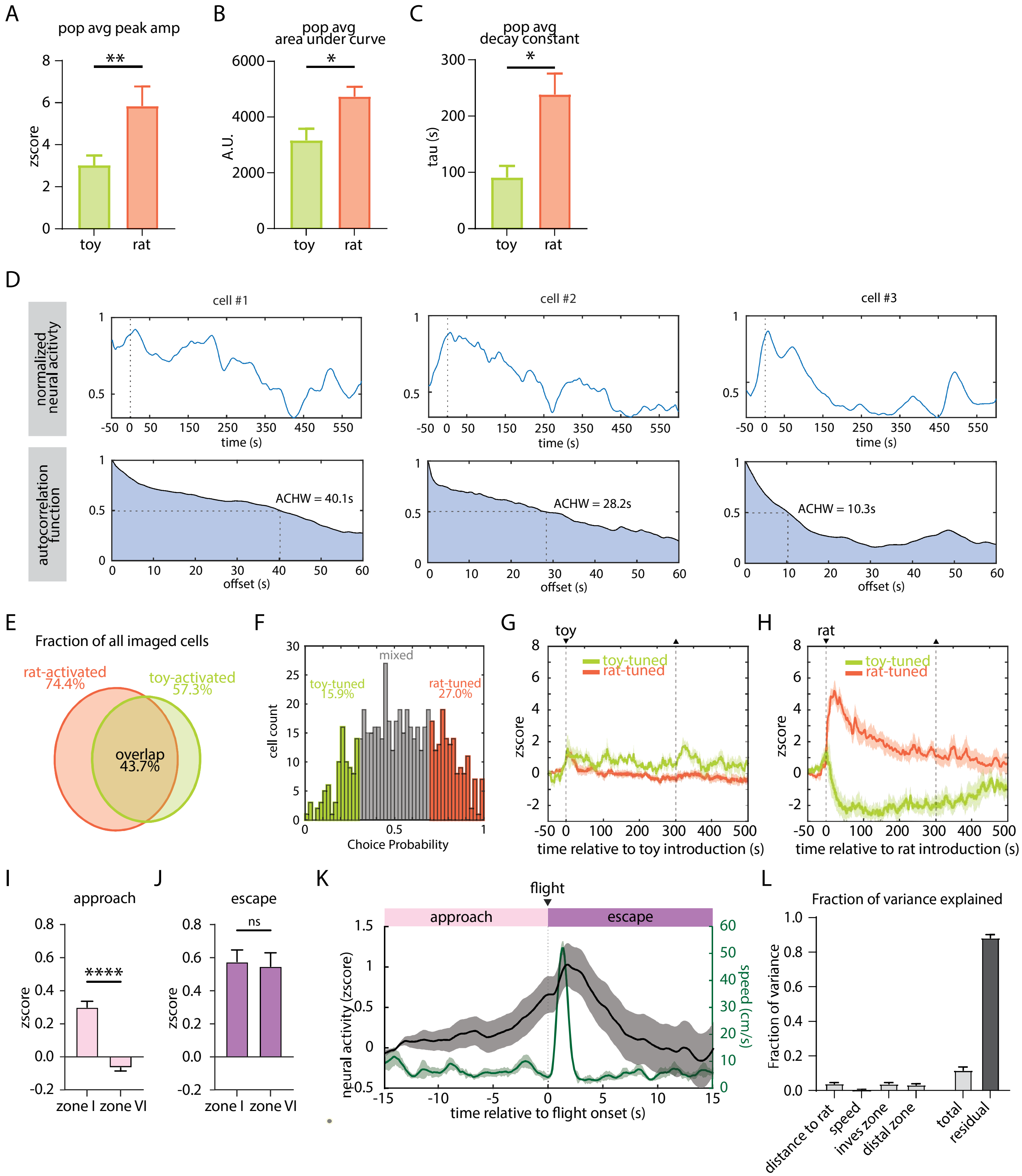

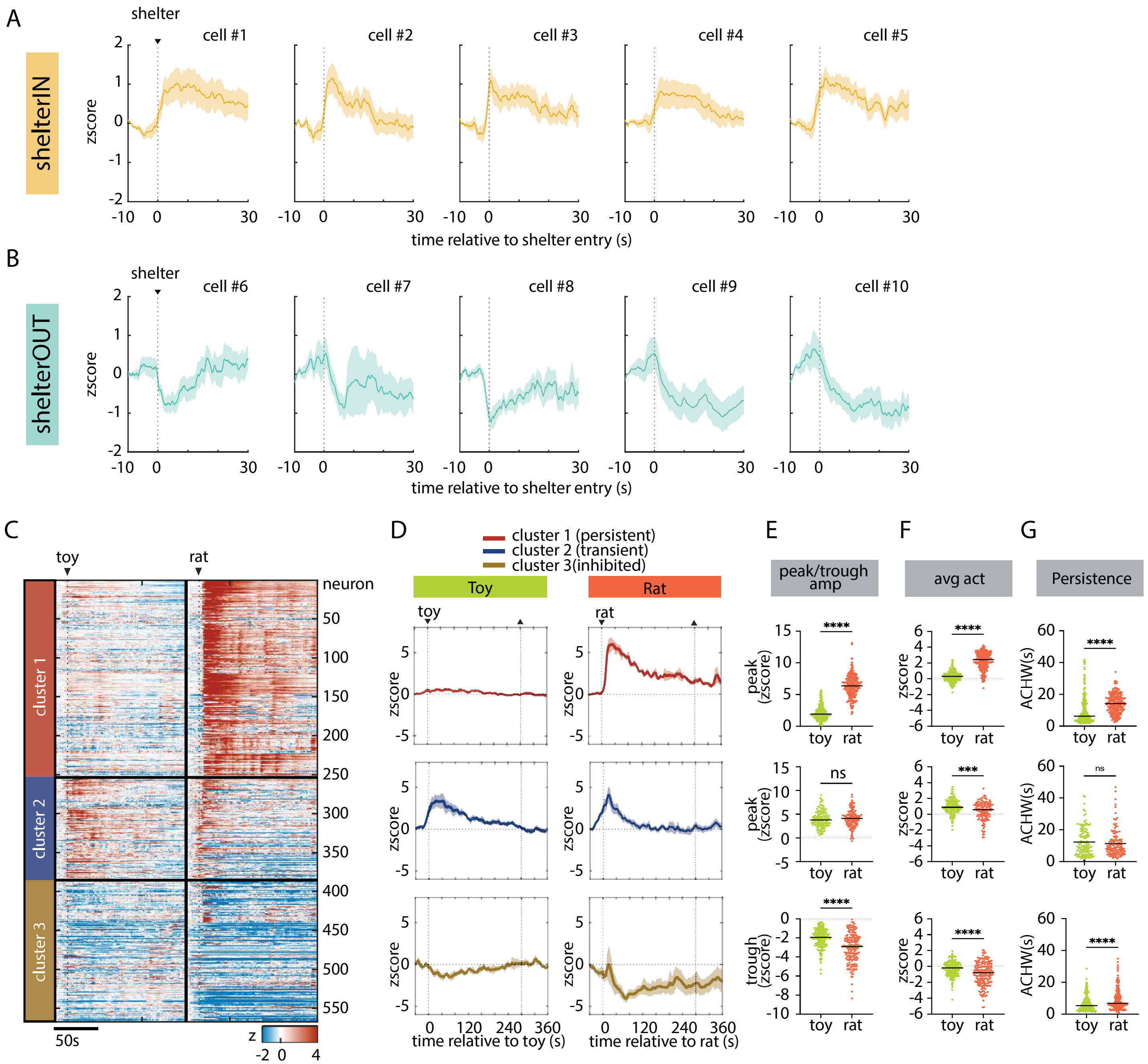

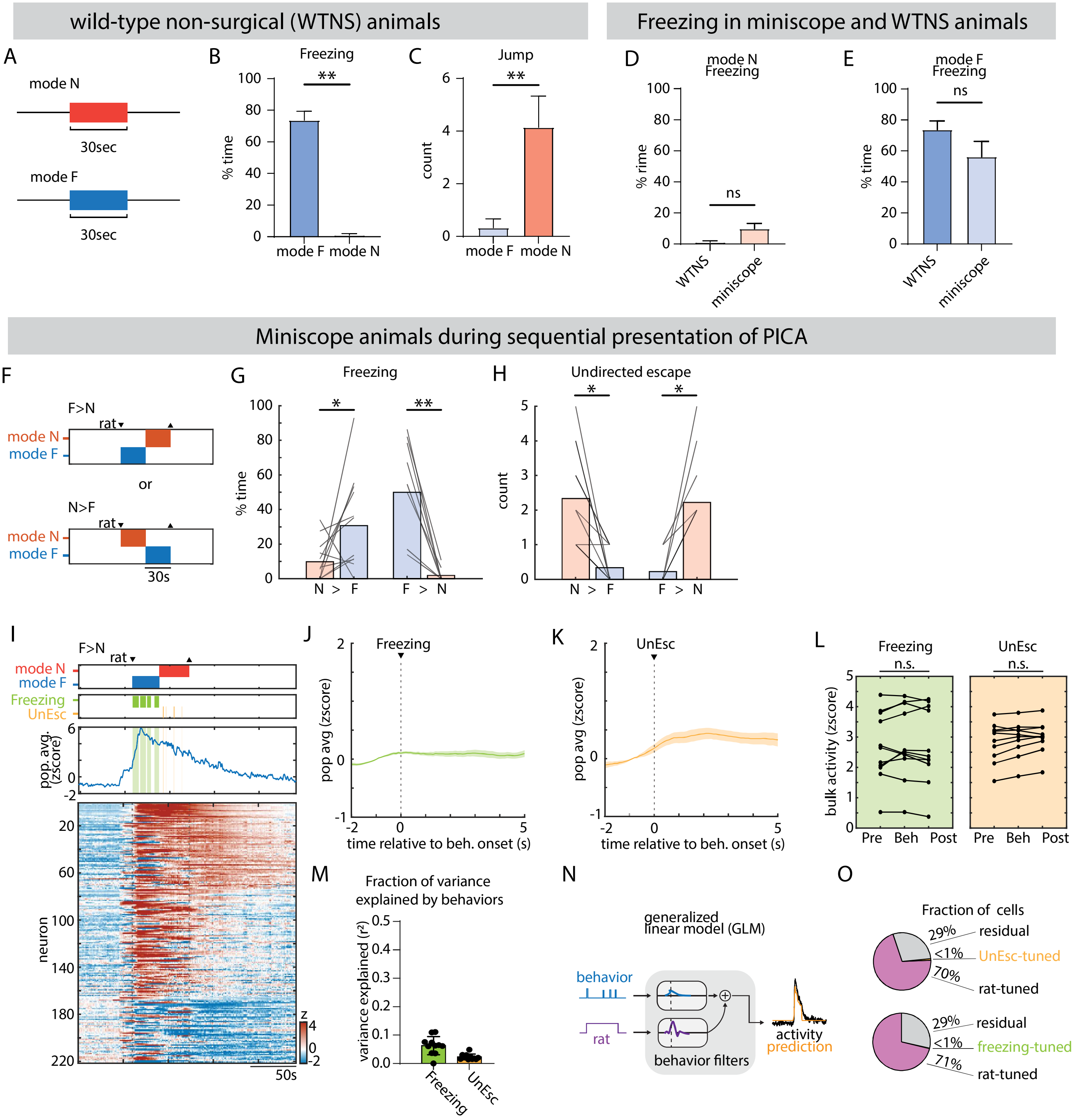

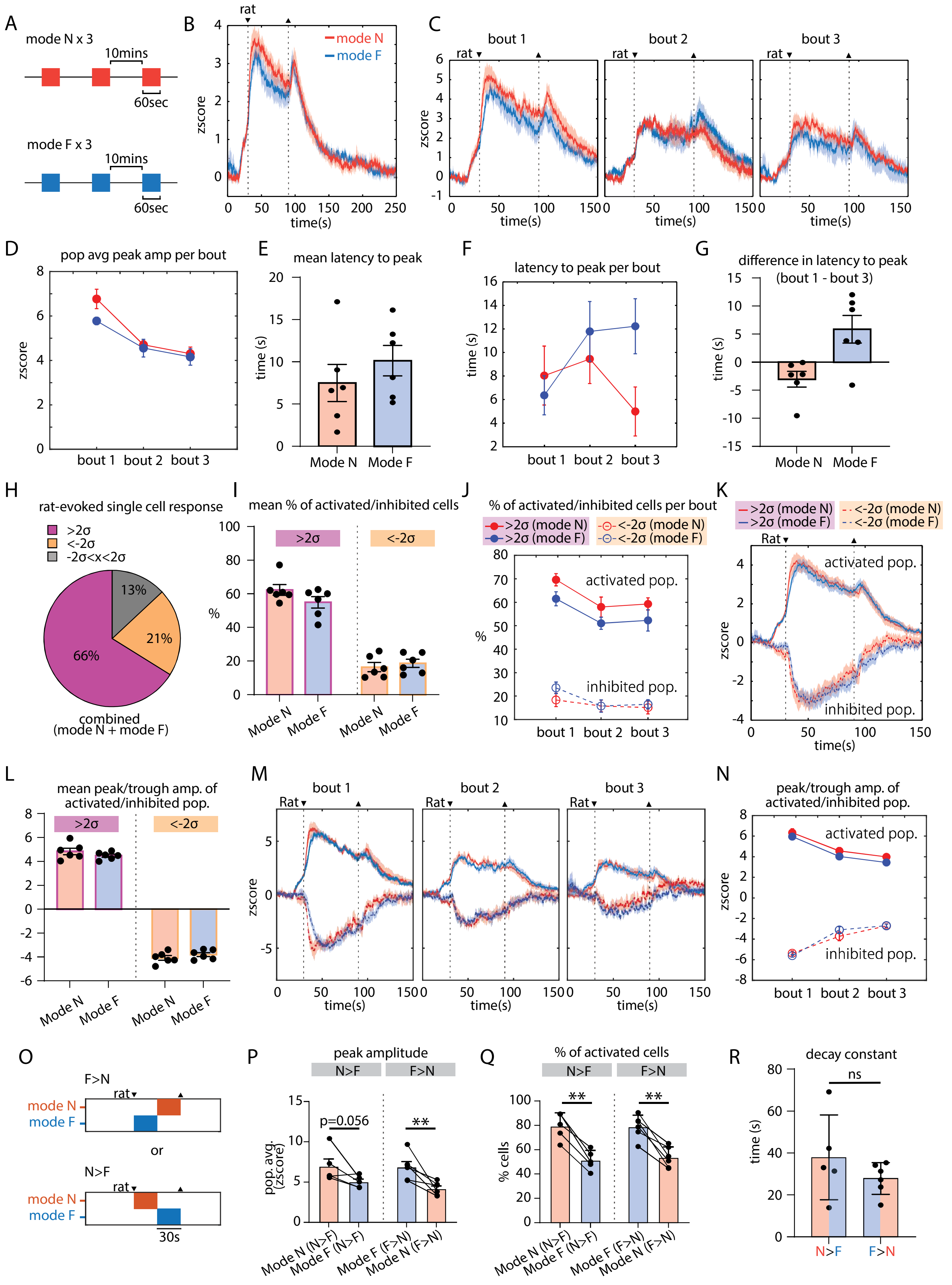

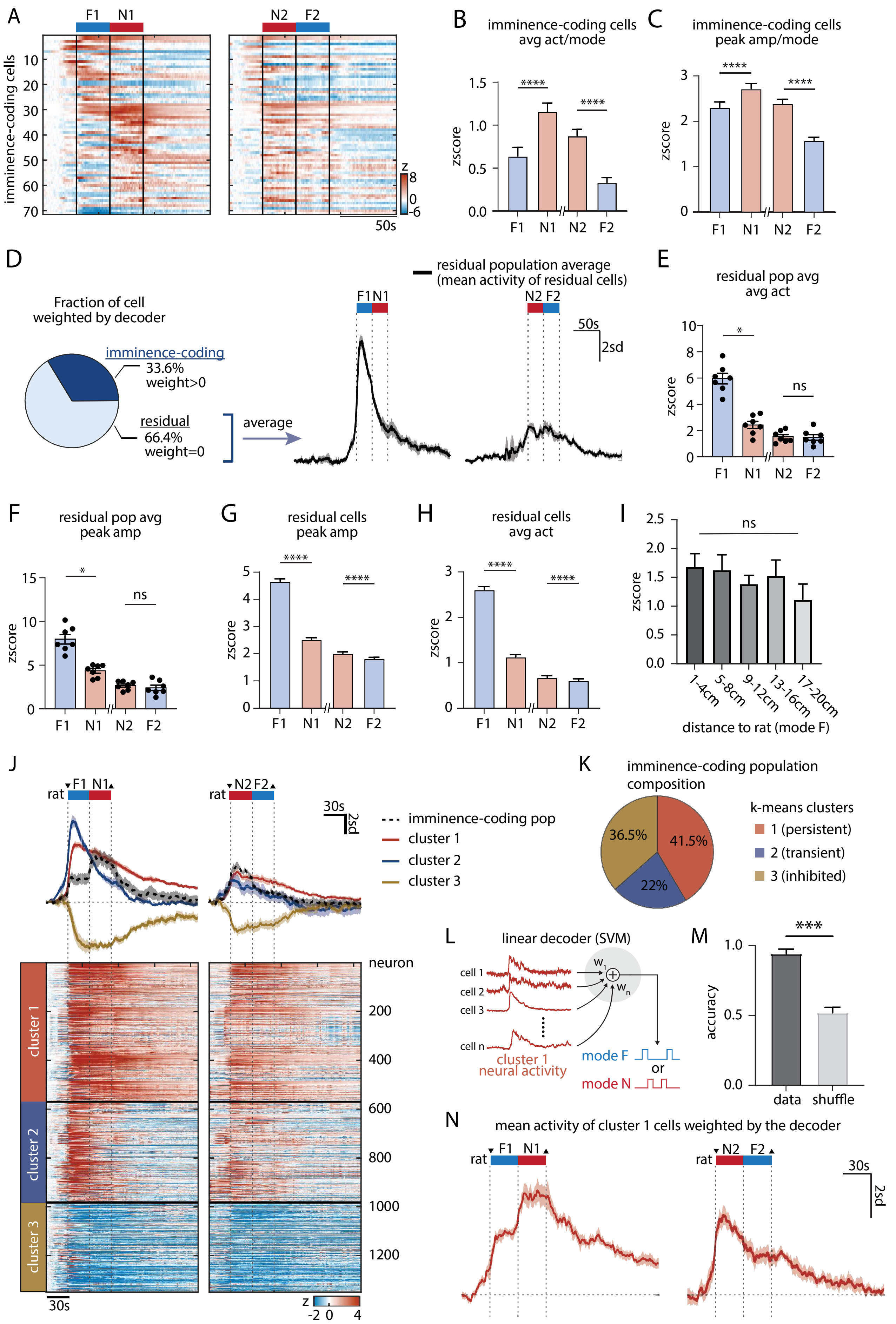

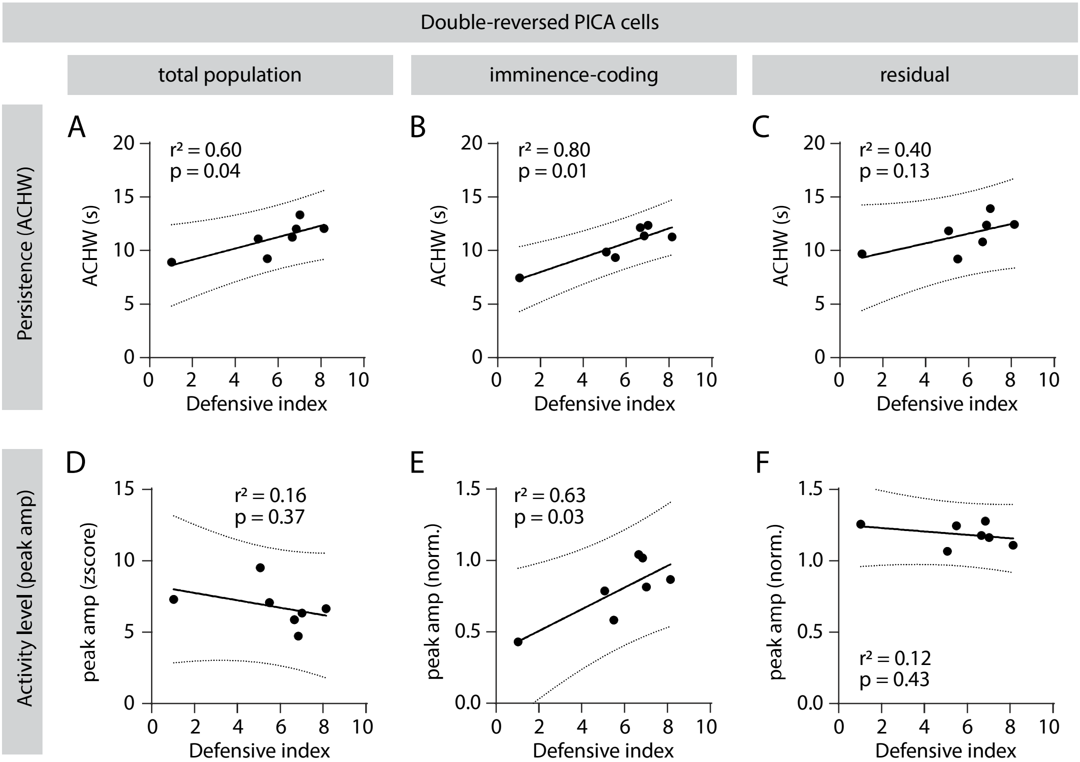

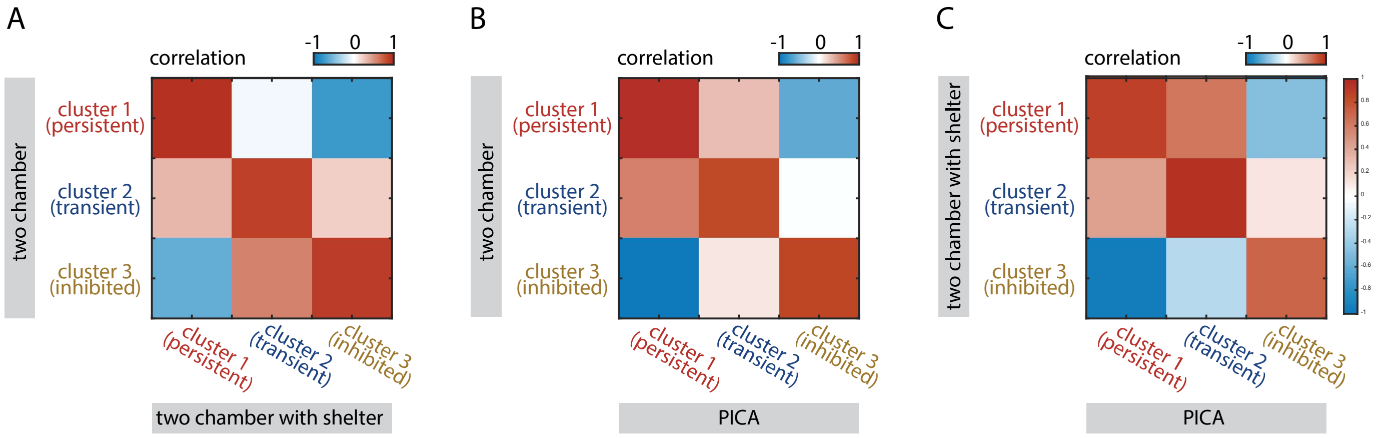
